# A co-conserved gene pair supports *Caulobacter* iron homeostasis during chelation stress

**DOI:** 10.1101/2024.10.16.618771

**Authors:** Sergio Hernandez Ortiz, Kiwon Ok, Thomas V. O’Halloran, Aretha Fiebig, Sean Crosson

## Abstract

Synthetic metal chelators are widely used in industrial, clinical, and agricultural settings, leading to their accumulation in the environment. We measured the growth of *Caulobacter crescentus*, a soil and aquatic bacterium, in the presence of the ubiquitous chelator ethylenediaminetetraacetic acid (EDTA) and found that it restricts growth by lowering intracellular iron levels. Using barcoded transposon sequencing, we identified an operonic gene pair, *cciT–cciO*, that is required to maintain iron homeostasis in laboratory media during EDTA challenge. *cciT* encodes one of four TonB-dependent transporters that are regulated by the ferric uptake repressor (Fur) and stands out among this group of genes in its ability to support *Caulobacter* growth across diverse media conditions. The function of CciT strictly requires *cciO*, which encodes a cytoplasmic Fe^II^ dioxygenase-family protein. Our results thus define a functional partnership between an outer membrane iron receptor and a cytoplasmic dioxygenase that are broadly co-conserved in Proteobacteria. We expanded our analysis to natural environments by examining the growth of mutant strains in freshwater from two lakes, each with biochemical and geochemical profiles that differ markedly from standard laboratory media. In lake water, *Caulobacter* growth did not require *cciT* or *cciO* and was less affected by EDTA treatment. This result aligns with our observation that EDTA toxicity is influenced by common forms of biologically chelated iron and the spectrum of free cations present in the medium. Our study defines a conserved iron acquisition system in Proteobacteria and bridges laboratory-based physiology studies with real-world conditions.

**IMPORTANCE:** Metal-chelating chemicals are widely used across industries, including as preservatives in the food sector, but their full impact on microbial physiology is not well understood. We identified two genes, *cciT* and *cciO*, that function together to support *Caulobacter crescentus* iron balance when cells are exposed to the common synthetic chelator, EDTA. CciT is an outer membrane transporter and CciO is a dioxygenase-family protein that are mutually conserved in many bacteria, including several human pathogens, where mutations in *cciT* homologs are linked to clinical resistance to the siderophore antibiotic, cefiderocol. This study identifies a conserved genetic system that supports iron homeostasis during chelation stress and illuminates the iron acquisition versatility and stress resilience of *Caulobacter* in freshwater environments.

## Introduction

Microbes are central to the productivity of aquatic and terrestrial ecosystems (1), playing a key role in the biogeochemical cycling of elements and the degradation of complex organic biopolymers into assimilable molecules (2). However, anthropogenic chemicals can perturb microbial physiology and community structure (3, 4) and affect ecological balance (5–9). The broad-spectrum metal chelator ethylenediaminetetraacetic acid (EDTA) is widely used in industrial applications (10) and in personal, medical, and food products (11, 12). Global production of EDTA and related synthetic chelators reaches hundreds of thousands of tons per year (13, 14). These chelators have been detected in the environment at varying concentrations and pose a potential threat to ecosystem health because they bind essential metal ions (10, 15), which can broadly perturb microbial physiology (16–18). Studies of microbial genes that influence fitness under chelation stress are needed as we work to understand the effects of these ubiquitous molecules on microbial biology. *Caulobacter* species provide excellent models for such studies, as they are ecologically (19) and metabolically (20–24) versatile and prevalent in both soil and aquatic ecosystems (19). These gram-negative bacteria have important roles in soils, where they are reported to enhance the growth of model plant systems (25, 26) and have been identified as ecological hub species in plant-associated microbial communities in the rhizosphere and phyllosphere (27, 28). Here we report a randomly barcoded transposon sequencing approach (RB-TnSeq) (29) to identify genes that contribute to *Caulobacter crescentus* fitness in a complex medium containing a growth-limiting concentration (300 µM) of EDTA. This approach led to the discovery of genes whose disruption diminished or enhanced bacterial fitness during EDTA challenge. Genes directly controlled by several metal-responsive regulators, including the ferric uptake repressor (Fur) (30, 31), the zinc uptake repressor (Zur) (32), and a uranium/zinc/copper-responsive two-component system (UzcRS/CusRS) (33, 34), were among those that influence fitness in the presence of EDTA. However, a subset of genes in the Fur regulon were identified as major fitness determinants at the tested growth-limiting EDTA concentration.

We focused specifically on a functional analysis of two genes that are directly regulated by Fur, are co-conserved and syntenic across Proteobacteria, and strongly support *C. crescentus* fitness in EDTA-containing media: *cciT*, encoding a TonB-dependent outer membrane transporter, and *cciO*, encoding an uncharacterized 2-oxoglutarate (2OG)/Fe^II^-dependent dioxygenase. CciT is related to the Fiu transporter of *E. coli* and the PiuA transporters of select gram-negative pathogens including *Acinetobacter baumannii, Burkholderia pseudomallei*, and *Pseudomonas aeruginosa*. Fiu and PiuA transporters import Fe^III^-catecholate siderophores, including the catecholate antibiotic cefiderocol (FDC) (35–38). Like *cciT, fiu* and *piuA* are typically adjacent to a 2OG/Fe^II^-dependent dioxygenase gene on the chromosome. To our knowledge, a functional connection between these syntenic genes has remained undefined to date.

In this study, we demonstrate that EDTA treatment results in decreased intracellular iron levels in *C. crescentus* and that the *cciT* and *cciO* genes are critical for growth under chelation stress. Through a systematic genetic analysis of Fur-regulated outer membrane transport systems, we show that the function of CciT has a strict – and specific – requirement for CciO. Our data thus demonstrate a central role for these two proteins in maintenance of iron homeostasis in the presence of a high-affinity chelator. Our study illuminates the versatility of *C. crescentus* iron-acquisition mechanisms in physicochemically-complex systems, advances understanding of the genetic foundations of bacterial resilience under chelation stress, and defines the functional roles of a broadly co-conserved pair of iron homeostasis genes in this context.

## Results

### Genetic contributors to *Caulobacter* fitness under EDTA chelation stress

To identify *Caulobacter* genes that affect fitness in the presence of EDTA, we implemented a randomly barcoded transposon (Tn) sequencing approach (RB-TnSeq) (29) using a previously described *C. crescentus Tn*-Himar mutant library (39, 40). We separately cultivated the library in liquid peptone-yeast extract (PYE) medium and in liquid PYE medium containing 300 µM EDTA. This concentration enabled the identification of genes with relative positive and negative effects on fitness in the presence of EDTA when they are disrupted, as it resulted in a 50% decrease in growth of the wildtype (WT) strain (Figure S1). The relative fitness of most mutant strains was unaffected by EDTA treatment (Figure 1A). Mutant strains with the lowest fitness scores in medium containing EDTA included those with defects in cell envelope and cell development functions (mutation of CCNA_01118, *bamE, fabZ, divJ*) or with predicted nutrient acquisition functions (mutation of CCNA_00028 and CCNA_03155-03159) (Figure 1A and Table S1). Mutants of the polar cell development regulator gene *divJ* (41) were among a group of several strains with fitness defects regardless of the presence or absence of EDTA in the medium (Table S1). While mutation of CCNA_01118, *bamE*, or *fabZ* led to basal fitness defects in untreated liquid medium, EDTA treatment exacerbated these defects. The CCNA_01118 gene is located adjacent to *divJ* and *fssA* (42), suggesting that it has a role in polar cell development. *fabZ* is part of a genetic locus with important functions in lipopolysaccharide and outer membrane biogenesis (43), while *bamE* encodes a component of the β-barrel protein assembly machine complex in *Caulobacter* (44). The EDTA sensitivity phenotype of *bamE* insertional mutants is consistent with a previous report (45). CCNA_00028 encodes a putative TonB-dependent transporter (TBDT) which belongs to a class of outer membrane proteins that function in the acquisition of vitamin B_12_, ferric siderophore complexes, nickel complexes, and carbohy-drates (46). Disrupting CCNA_00028 had no effect on fitness in liquid medium without EDTA but led to lower fitness in the presence of EDTA. In fact, mutants of CCNA_00028 had the most severe fitness defects in EDTA, relative to untreated medium, in the entire dataset (Table S1). Other mutants with large relative defects in the presence of EDTA included mutants in the directly adjacent gene CCNA_00027, encoding a likely 2OG/Fe^II^-dependent dioxygenase (Figure 1B), and in CCNA_03155–03159, encoding a PepSY-like system implicated in periplasmic reduction of ferric iron and iron import (47–50) (Table S1 and Figure 1A). CCNA_00028, CCNA_00027, and the CCNA_03155–03159 genetic locus are direct regulatory targets of the ferric uptake repressor Fur (30, 31), suggesting a connection between iron homeo-stasis and EDTA resistance at the tested concentration (300 µM).

**Figure 1.**
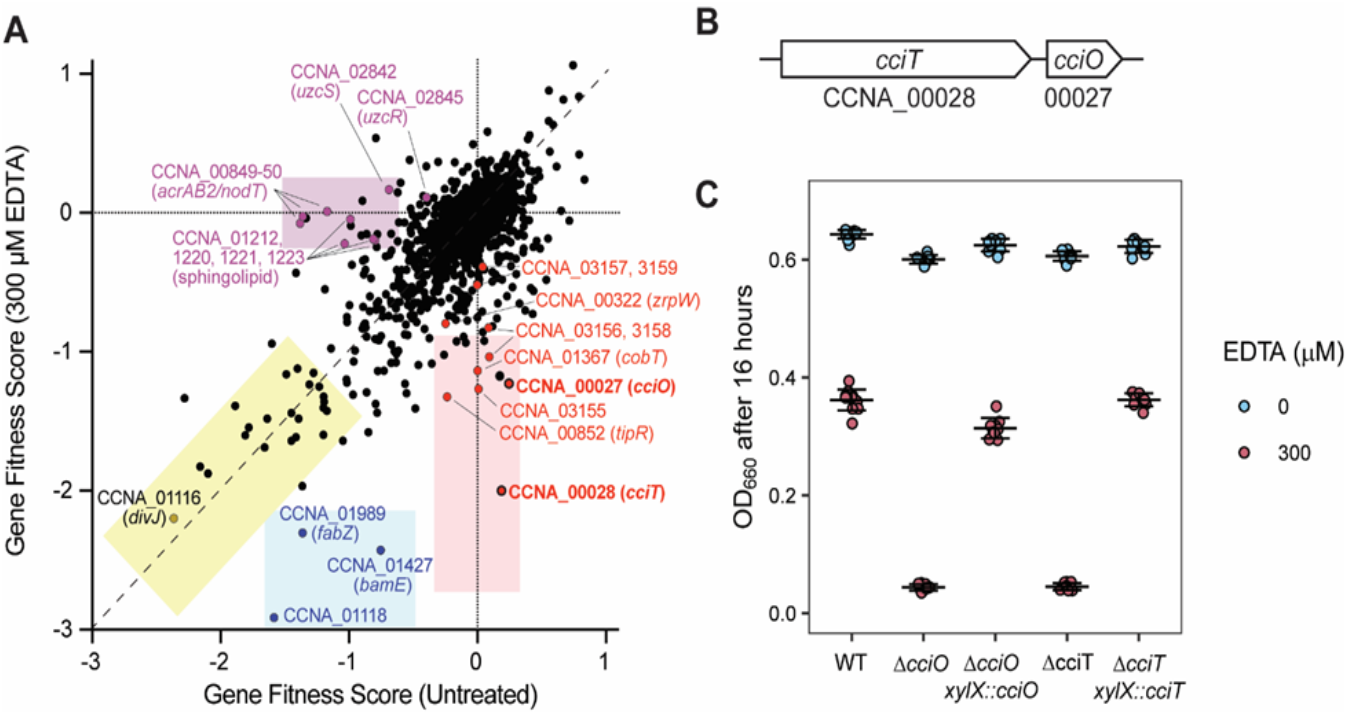
Identification of genes that contribute to *C. crescentus* fitness in the presence of EDTA. **(A**) Fitness scores of C. crescentus genes after cultivation of a barcoded Tn-himar mutant library in complex PYE medium (Untreated) or in the same medium containing 300 μM EDTA for 16 h. Each point represents one gene. Scores represent the average of four independent cultures in each condition. Multiple classes of fitness defects are observed: the yellow box highlights genes that lead to similar growth defects when disrupted regardless of EDTA; the blue box shows genes that have a fitness defect in PYE medium when disrupted that is more severe in the presence of EDTA; the pink box highlights genes that have a fitness defect when disrupted specifically in PYE medium containing EDTA; the purple box highlights genes that have a basal fitness defect in PYE medium that is rescued by the addition of EDTA. Representative genes for each class are shown. The complete fitness data set is presented in Table S1. **(B)** Diagram of the cciT–cciO locus. **(C)** Optical density at 660 nm of the wild-type (WT) strain and strains bearing in-frame deletions of cciO (DcciO) or cciT (DcciT), and deletion strains expressing the gene from its native promoter from an ectopic insertion site after 16 h of growth in liquid PYE medium alone (blue) or containing 300 μM EDTA (red). All cultures were inoculated at a starting optical density (OD_660_) of 0.01. Box plots reflecting the median and the 25th and 75th percentiles are overlaid with the individual data points for each independent culture (WT, n = 24; ΔcciO and ΔcciT, n = 18; complementation strains, n = 6).

Phenotypic effects of mutations in several zinc uptake regulator (Zur)-regulated genes have previously been reported in *C. crescentus* at higher EDTA concentrations (500 µM and 800 µM) on solid M2 defined medium (32). However, disrupting genes in the Zur regulon (32) individually did not have noteworthy fitness defects in our EDTA Tn-seq experiment except for *zrpW* (CCNA_00322), which encodes an uncharacterized transmembrane protein: *zrpW* mutants showed a small (factor ∼1.5) reduction in fitness in the presence of 300 µM EDTA (Table S1; see Materials and Methods for fitness calculation method). Disruption of genes reported to support fitness under copper-induced stress (51–53) also did not strongly affect fitness except for *cobT* (CCNA_01367), encoding a predicted component of the cobaltochelatase complex involved in vitamin B_12_ biosynthesis. Strains harboring transposon insertions in the uranium/zinc/copper-responsive regulatory system (*uzcRS;* homologous to *E. coli cusRS*) (33) had a basal fitness defect in liquid medium alone that was rescued by treatment with EDTA. The *acrAB2*/*nodT* antibiotic resistance locus (54) is directly activated by the UzcRS two-component system (33), and insertional mutants in *acrAB2*/*nodT* had the largest EDTA rescue phenotype of all *C. crescentus* mutants, as indicated by their low fitness score in the absence of EDTA and their normal score in the presence of EDTA (Figure 1A and Table S1). In contrast, strains with insertions in *tipR*, the transcriptional repressor of *acrAB2/nodT* (55, 56), displayed the opposite fitness profile, consistent with the known regulatory connection between these genes. The basal fitness defects of strains with insertions in sphingolipid biosynthesis genes (CCNA_01212, CCNA_01220, CCNA_01221, CCNA_01223) (57) were also rescued by the addition of EDTA to the growth medium. We conclude that genes with cell envelope and iron acquisition functions are key *Caulobacter* fitness determinants in the presence of 300 µM EDTA. Our observation that EDTA treatment rescues the mutant phenotypes of select cell envelope mutants suggests a complex interaction between metal ion homeostasis and cell envelope development mechanisms in *C. crescentus*.

### CCNA_00027 (*cciO*) and CCNA_00028 (*cciT*) are co-conserved EDTA resistance genes

Given the strong and specific fitness defects seen for mutants of the two genes encoding a TBDT (CCNA_00028) and dioxygenase (CCNA_00027) in complex medium containing EDTA, we chose these genes for further study. There was no obvious functional relationship between these two gene families: TBDTs transport a diverse array of molecules across the outer membrane, while 2OG/Fe^II^-dependent dioxygenases introduce diatomic oxygen into a range of substrates (58). However, both genes are directly regulated by Fur (30, 31) and are operonic (59), suggesting related function(s).

RB-TnSeq data (Table S1) could not differentiate whether the fitness defect ascribed to disruption of the TBDT gene (CCNA_00028) was due to a polar effect on the adjacent dioxygenase gene (CCNA_00027) or to the disruption of CCNA_00028 *per se* (see Figure 1B). To distinguish between these two possibilities and to validate the RB-TnSeq results, we constructed mutants carrying in-frame, unmarked deletions of CCNA_00027 or CCNA_00028. We hereafter refer to these genes as *cciO* and *cciT*, respectively, for reasons we will explain below. Both in-frame deletion mutants grew like the WT strain in complex medium and failed to grow in the same medium with 300 µM EDTA added (Figure 1C), confirming the RB-TnSeq fitness measurements (Figure 1A; Table S1). The growth defect of each mutant in complex liquid medium containing EDTA was genetically complemented by ectopic expression of the deleted gene from its native promoter (Figure 1C), thus defining CciO and CciT as EDTA resistance factors.

We assessed the prevalence and conservation of *cciO* and *cciT* through a pangenome analysis that binned genes into orthologous clusters (60). A comparison of 69 *Caulobacter* isolates from diverse environments including water, soil, and plant-associated ecosystems identified a highly related cluster of dioxygenase genes that includes *cciO*, which was detected in all but one of the analyzed genomes. The TBDT gene cluster that included *cciT* was present in 80% of the analyzed genomes (Table S2). A gene neighborhood analysis (61) showed that *cciO* homologs are commonly located directly adjacent to *cciT* homologs in *Caulobacter* species and more broadly in the Proteobacteria (Figure 2). This high degree of synteny supports an evolutionary and functional relationship between *cciT* and *cciO*.

**Figure 2.**
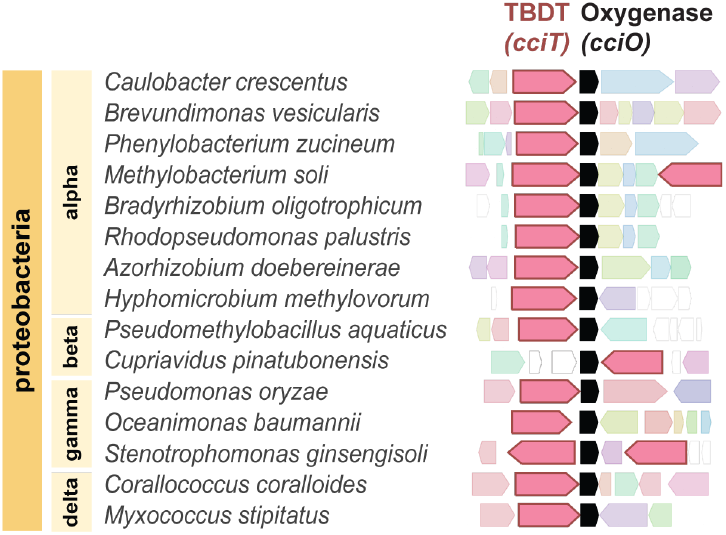
Orthologs of *cciT* and *cciO* are syntenic in Proteobacteria. Gene neighborhood analysis anchored on cciO reveals co-inheritance of cciO and cciT orthologs. Neighborhood configuration was defined by selecting the highest-scoring BLAST hits from different proteobacterial groups and identifying the corresponding neighborhoods using webFLAGs (61). Representative genomes across the phylum Proteobacteria are presented. The cciT and cciO orthologs are colored in pink and black, respectively.

### Growth defects of *cciO* and *cciT* mutants are rescued by chelated iron

Having identified *cciT* and *cciO* as EDTA resistance genes, we sought to define the functions of the proteins encoded by this co-conserved gene pair. The growth of Δ*cciO* and Δ*cciT* mutants in complex (PYE) liquid medium was equivalent to that of the WT strain (Figure S2A & Figure 1C). When we spotted a ten-fold dilution series on solid complex (PYE) medium, we observed comparable numbers of colony-forming units (CFUs) for all strains, although Δ*cciO* and Δ*cciT* strains produced visibly smaller colonies (Figure 3A). The colony size defect was genetically complemented by expressing each of the deleted genes from an ectopic locus from its own promoter (Figure S3A). The fact that transcription of *cciO* and *cciT* is regulated by Fur (30, 31) indicated that the diminished growth on solid medium seen in their respective mutants might be a result of a defect in iron homeostasis. To test this hypothesis, we added 10 µM Fe•EDTA chelate to solid PYE agar medium, which restored the colony size of the mutants to WT levels (Figure 3B). It was possible that slower growth impacted cell size. We used light microscopy to measure the cell sizes of these strains when grown in PYE broth and found that the Δ*cciO* and Δ*cciT* mutant cells were ∼25% smaller than WT (p<0.0001) (Figure 3E & Figure S3B).

**Figure 3.**
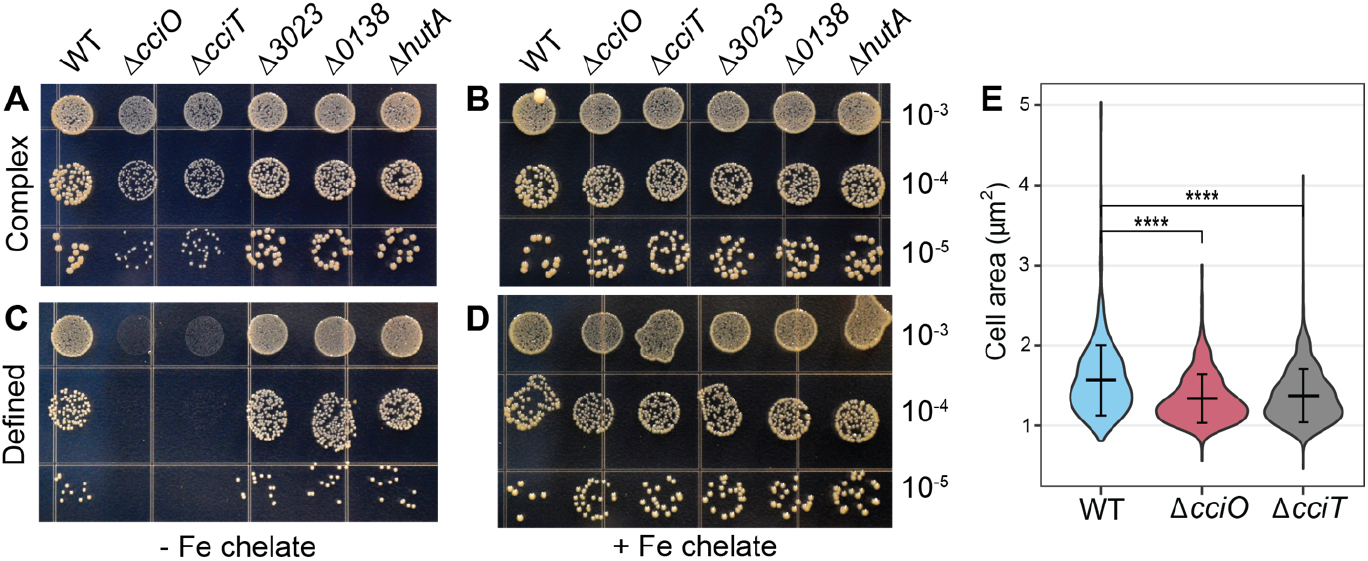
*cciT* and *cciO* deletion mutants have colony size and cell size phenotypes. Representative photographs of growth on plates for the WT strain, the ΔcciO deletion mutant, and deletion mutants for each of the Fur-regulated TonB-dependent transporter genes (ΔcciT, ΔCCNA_00138, ΔCCNA_03023, and ΔhutA) serially diluted and spotted onto complex (PYE**; A & B**) or defined (M2X**; C & D**) agar medium. The medium in **B** and **D** was supplemented with 10 μM Fe•EDTA chelate, whereas the medium in **A** and **C** was not. The dilution factor of each row is indicated at right. Panel **E** shows the distribution of cell areas in WT (n=759), ΔcciO (n=750), and ΔcciT (n=1154) cells grown in PYE broth for 6 hours. Comparisons between strains were made using a Kruskal-Wallis test, followed by Dunn’s post test. **** p < 0.0001

We tested strain growth in liquid M2X defined medium containing 10 µM Fe•EDTA as an iron source. The growth of the Δ*cciO* and Δ*cciT* mutants matched that of the WT strain in this condition (Figure S2B). Replacing chelated iron (as Fe•EDTA) with 10 µM FeCl_3_ did not affect WT growth but resulted in attenuated growth of the Δ*cciO* and Δ*cciT* mutants (Figure S2B). We conclude that *cciO* and *cciT* mutants have a decreased ability to acquire unchelated Fe^III^ in defined liquid medium. On defined solid (agar) medium without added iron, the WT *C. crescentus* strain had sufficient iron scavenging ability to grow well (Figure 3C), whereas growth of the Δ*cciO* and Δ*cciT* mutants was severely compromised. Again, this defect was rescued by the addition of 10 µM Fe•EDTA chelate (Figure 3D), indicating that both *cciO* and *cciT* support acquisition of iron from a defined solid medium. Accordingly, we named the constituent genes of this syntenic gene pair *<u>C</u>. crescentus* iron transporter (CCNA_00028; *cciT*) and *<u>C</u>. crescentus* iron oxygenase (CCNA_00027; *cciO*); we provide additional experimental data supporting a functional role for *cciT* and *cciO* in cellular iron homeostasis below.

### Deletion of *cciT* or *cciO* de-represses transcription of the Fur regulon

To test the hypothesis that intracellular iron homeostasis is perturbed in Δ*cciT* and Δ*cciO*, we first measured global transcript levels by transcriptome deep sequencing (RNA-seq), reasoning that lower intracellular iron levels would manifest as de-repressed transcription of Fur-regulated genes. Deletion of *cciT* or *cciO* resulted in increased expression of a small and highly congruent gene set (Figure 4 and Table S3). Of the 24 genes differentially expressed in at least one of the mutants (absolute log_2_[fold change] > 1; FDR *p*-value < 10^™4^), 17 had increased transcript levels in both mutants, and 21 had been demonstrated to be regulated in an iron-dependent manner in earlier studies (30, 62). Most genes with enhanced expression in Δ*cciT* and Δ*cciO* are known to be repressed by Fur (30, 31, 62), including *cciT* itself, three additional TBDT genes, and the predicted inner membrane ferrous iron transport system *feoAB*. To validate the RNA-seq results, we measured transcription using reporter constructs consisting of the fluorescent reporter gene *mNeonGreen* cloned downstream of the *cciT* or *feoAB* promoter. Consistent with the RNA-seq data, transcription from the *cciT* and *feoAB* promoter constructs was significantly de-repressed in Δ*cciT* and Δ*cciO* relative to WT (Figure S4A & B; Figure 4C).

**Figure 4.**
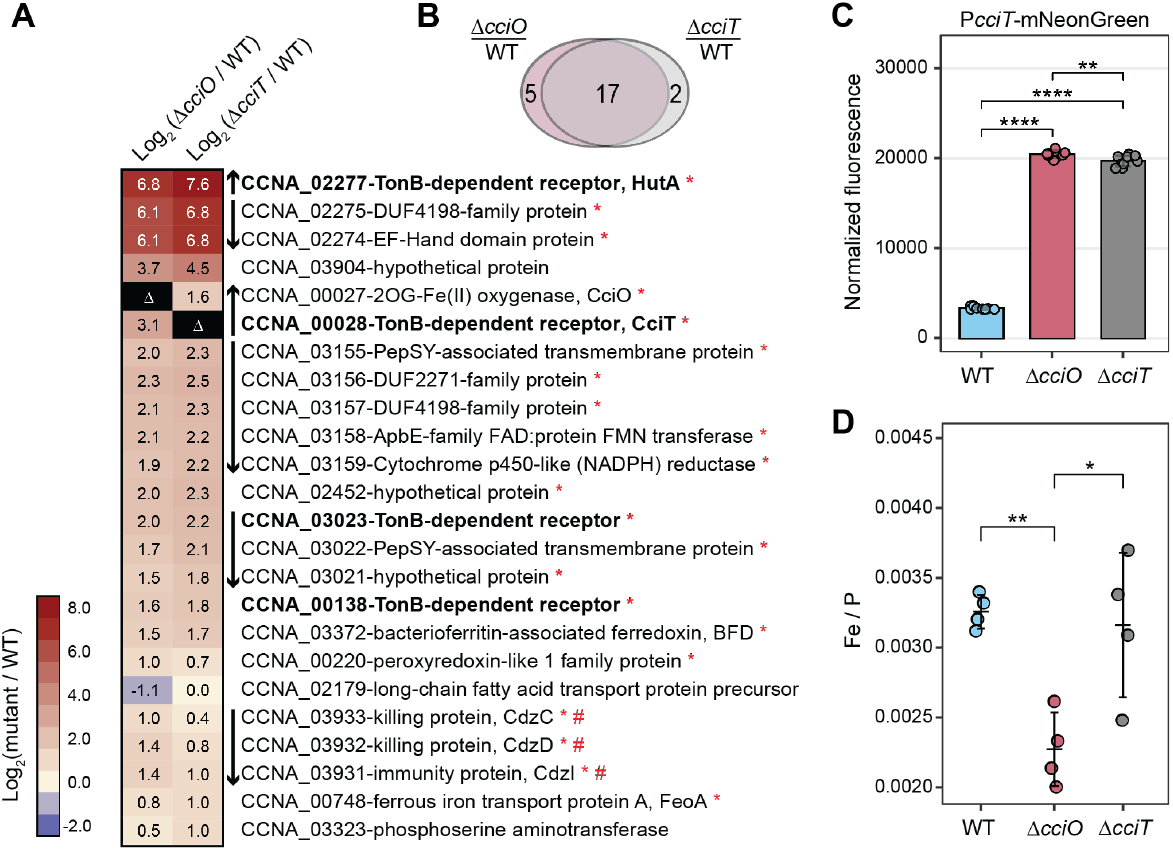
Δ*cciT* and Δ*cciO* mutants have an iron homeostasis defect. **(A)** Heatmap of expression of the 24 genes differentially regulated in ΔcciT or ΔcciO strains, relative to WT cells, grown in defined medium containing FeCl_3_ as a sole iron source. Each row represents one gene, each column represents a mutant compared to the WT culture, and the color of each cell reflects the magnitude of change, as the log_2_(mutant/WT) expression level (see key at left). Boldface indicates the TBDT genes directly repressed by Fur and upregulated during iron limitation. Vertical arrows between the heatmap and gene listings indicate the orientations of adjacent genes (presumed operon structures). Red asterisks indicate genes with a Fur-binding motif in their promoters (31). Differentially expressed genes in the mutants compared to the WT strain were defined as those with an absolute log_2_(fold change) > 1 and a false discovery rate (FDR) p-value < 0.0001. **(B)** Venn diagram showing the extent of overlap between differentially expressed genes in each mutant compared to the WT strain. **(C)** Transcription from the cciT promoter (P_cciT_) using a P_cciT_:mNeonGreen reporter plasmid in WT, DcciT, and DcciO strains grown in PYE complex medium (n = 9). Comparisons of normalized fluorescence were made with a Welch’s ANOVA test followed by Tukey’s multiple comparison test. Statistical significance was indicated as follows: **, p < 0.01; ****, p < 0.0001. **(D)** Inductively coupled plasma mass spectrometry (ICP-MS) quantification of total cellular ^56^Fe, normalized to cellular phosphorus, in WT, ΔcciT, and ΔcciO strains grown in PYE complex medium. Comparisons of ^56^Fe levels between mutant strains and the WT were performed using ANOVA, followed by Tukey’s multiple comparison test. Statistical significance is indicated as follows: *, p < 0.05 ; **, <p < 0.01.

Additionally, we tested whether increased levels of *cciT* transcripts in Δ*cciO* led to increased CciT protein abundance in the outer membrane. Indeed, we detected significantly higher CciT protein levels in purified membranes of Δ*cciO* relative to the WT strain (Figure S4C & D) during iron limitation imposed by treatment with the divalent metal chelator dipyridyl. This result supports the RNA-seq data and demonstrates that Δ*cciO* phenotypes are not due to diminished CciT production.

### *cciT* and *cciO* mutants are resistant to iron-dependent antimicrobial compounds

To further assess the consequences of *cciT* or *cciO* deletion on cellular iron homeostasis, we used a streptonigrin sensitivity assay. Streptonigrin toxicity increases with increasing intracellular iron levels (63, 64), and both Δ*cciO* and Δ*cciT* strains were less sensitive than WT cells to streptonigrin when grown on PYE agar (Figure 5A & 5C), supporting the hypothesis that intracellular iron concentrations are lower in these strains. Supplementing the medium with 10 µM Fe•EDTA returned mutant sensitivity to streptonigrin to WT levels (Figure 5A & 5C). As a complementary approach, we measured the sensitivity of WT, Δ*cciO*, and Δ*cciT* strains to the drug CHIR-090, which inhibits the lipid A biosynthesis enzyme, LpxC (65). Lipid A is typically an essential component of the Gram-negative outer membrane, but the de-repression of Fur-regulated transcription (which occurs during iron limitation) renders lipid A dispensable in *C. crescentus* (57). CHIR-090 toxicity is therefore predicted to be diminished when cells are iron-limited. Consistent with this hypothesis, the *ΔcciO* and *ΔcciT* mutants were both more resistant to CHIR-090 treatment than the WT. Again, addition of 10 µM Fe•EDTA to the medium sensitized the mutants to this antimicrobial compound (Figure 5B & 5C). These data provide independent, pharmacological lines of evidence that cells lacking CciO or CciT have lower intracellular iron levels than WT.

**Figure 5.**
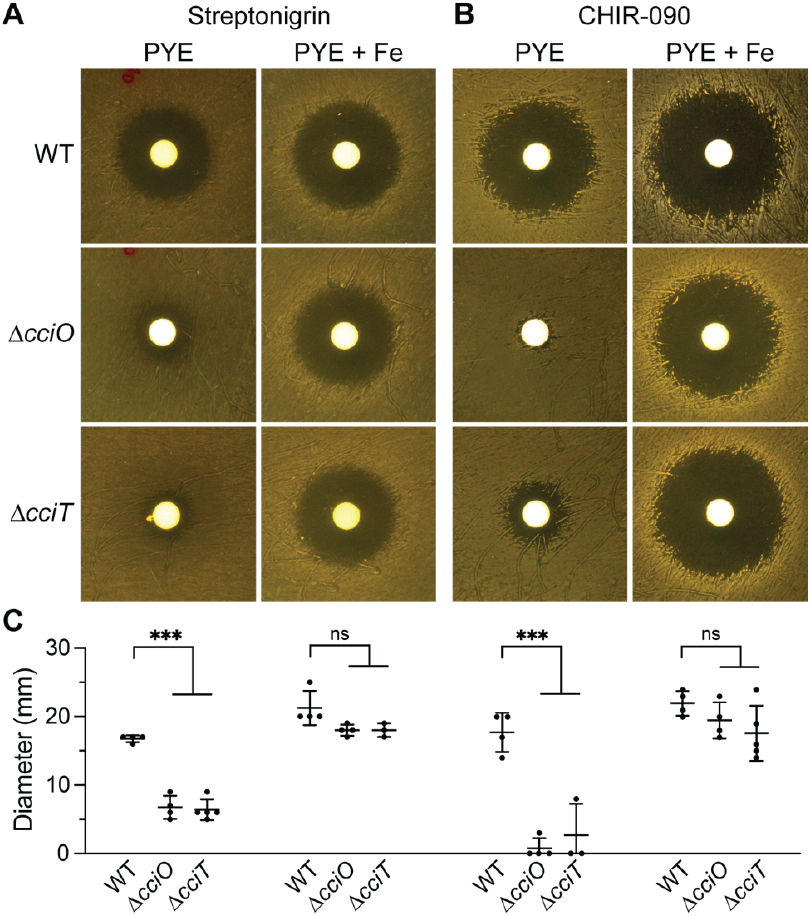
Δ*cciO* and Δ*cciT* mutants are resistant to streptonigrin and CHIR-090. WT, Δ*cciO*, Δ*cciT* were spread onto solid PYE medium without or with 10 mM Fe•EDTA and challenged with diffusion disks loaded with (**A**) streptonigrin or (**B**) CHIR-090. **(C)** Diameters of the zones of inhibition after growth for the plates shown in (A) and (B). The mean ± standard deviation of at least three independent experiments is overlayed with the individual data points. For each growth condition, the inhibitory zones were compared using analysis of variance (ANOVA) followed by a Dunnett’s post-hoc test. not significant, ns, p > 0.01; *** p < 0.001.

### Direct measurement of cellular iron levels in *cciT & cciO* mutants

The transcriptomic (Figure 4A) and pharmacological (Figure 5) experiments described above provide three independent lines of evidence that intracellular iron levels are lower in Δ*cciT* and Δ*cciO* cells. To directly measure the effect of *cciT* or *cciO* deletion on cellular iron levels, we turned to inductively coupled plasma triple-quadrupole mass spectrometry (ICP-MS-QQQ). Cells collected from plates contained contaminants from the agar, so these experiments were conducted on broth-grown cells. While Δ*cciT* and Δ*cciO* cells exhibited highly diminished growth and de-repressed transcription of Fur-regulated genes in a defined medium supplemented with FeCl_3_, ferric oxides in the cell pellets confounded our cellular iron measurements in this condition. Therefore, we conducted measurements on cells grown in PYE broth. As several biologically-relevant divalent cations are strongly chelated by EDTA, we tested whether iron, copper and zinc levels were basally affected in the Δ*cciT* and Δ*cciO* mutants cultivated in a complex medium. Metal abundance data were first normalized to colony forming units (atoms/CFU); copper and zinc showed no difference between WT and mutant strains while cellular iron levels appeared lower in Δ*cciO* (Figure S5). To more accurately assess differences in iron between strains, we further normalized the element concentration data to the levels of phosphorus in each sample, which is an established internal standard (66). Phos-phorus-normalized iron levels were significantly lower in Δ*cciO* (*p* < 0.007), with approximately 30% less iron than WT cells (Figure 4D). Δ*cciT* normalized iron levels were more variable than either WT or Δ*cciO* but did not significantly differ from the WT strain. This result, when considered alongside the transcriptomic data (Figure 4A) and pharmacological data (Figure 5), supports a model in which Δ*cciT* cells can bind iron but not efficiently import it. We discuss this model below.

### CciT and CciO coordinately function as a major iron-acquisition system in complex and defined media

Deletion of *cciT* or *cciO* decreased intracellular iron levels, but *cciT* is only one of four Fur-regulated TBDT genes that are transcriptionally activated in response to iron limitation in complex medium (30). We therefore tested whether and how the remaining Fur-regulated TBDT genes might contribute to *C. crescentus* fitness under our assay conditions. To this end, we generated mutant strains lacking each of the other three Fur-regulated TBDT genes: CCNA_00138, CCNA_03023, and CCNA_02277 (previously named *hutA* (67)). Growth of these single TBDT mutants was equivalent to that of WT cells on solid complex or solid defined medium, regardless of iron supplementation (Figure 3). Thus, the Δ*cciT* mutant was the only mutant in a Fur-regulated TBDT gene to exhibit a growth defect on solid medium.

To more rigorously define the contributions of the four Fur-regulated TBDTs to *C. crescentus* growth and to test the hypothesis that CciT acts as the main outer membrane iron importer in both complex PYE and defined M2X media, we developed a new set of mutant strains with deletions in three of the four Fur-regulated TBDT genes, leaving only one Fur-regulated TBDT gene intact in each strain (ΔΔΔ*cciT*^+^, ΔΔΔ*138*^+^, ΔΔΔ*3023*^+^, and ΔΔΔ*hutA*^+^). We also generated a quadruple deletion strain lacking all four Fur-regulated TBDT genes (ΔΔΔΔ). All strains had similar growth kinetics in liquid PYE medium during the exponential phase, indicating that *C. crescentus* can acquire iron through other transport systems in complex medium (Figure S2C). On solid complex medium, all triple deletion strains retaining only one active single transporter yielded comparable CFU numbers across serial dilutions, although only the ΔΔΔ*cciT*^+^ triple mutant had a colony size similar to that of WT cells (Figure 6A). The growth benefit provided by *cciT* was particularly evident on solid M2 defined medium without added iron, where the growth of strains ΔΔΔ*138*^+^, ΔΔΔ*3023*^+^, and ΔΔΔ*hutA*^+^ was diminished while that of ΔΔΔ*cciT*^+^ matched the growth of the WT strain (Figure 6C). Growth of the quadruple TBDT gene deletion strain (ΔΔΔΔ) was severely compromised relative to the WT strain on solid complex and defined media (Figure 6A & C). We conclude that at least one of the four Fur-regulated TBDT genes is required for growth on solid medium, and that CciT alone is sufficient to confer WT-like growth. Supplementation of M2X solid medium with 10 µM Fe•EDTA chelate completely rescued the growth of all strains except the ΔΔΔΔ strain which was only partially rescued by this concentration of EDTA-chelated iron (Figure 6B & D). This suggests that *C. crescentus* can likely acquire iron from Fe•EDTA through a system distinct from the four TBDTs studied here, although the limited growth observed in the ΔΔΔΔ strain may be partially supported by iron chelators present in agar. Robust chemical rescue of the strains retaining a single Fur-regulated TBDT gene by 10 µM Fe•EDTA provides evidence that each of these TBDTs can support acquisition of iron from Fe•EDTA at a concentration of 10 µM (Figure 6B & D).

**Figure 6.**
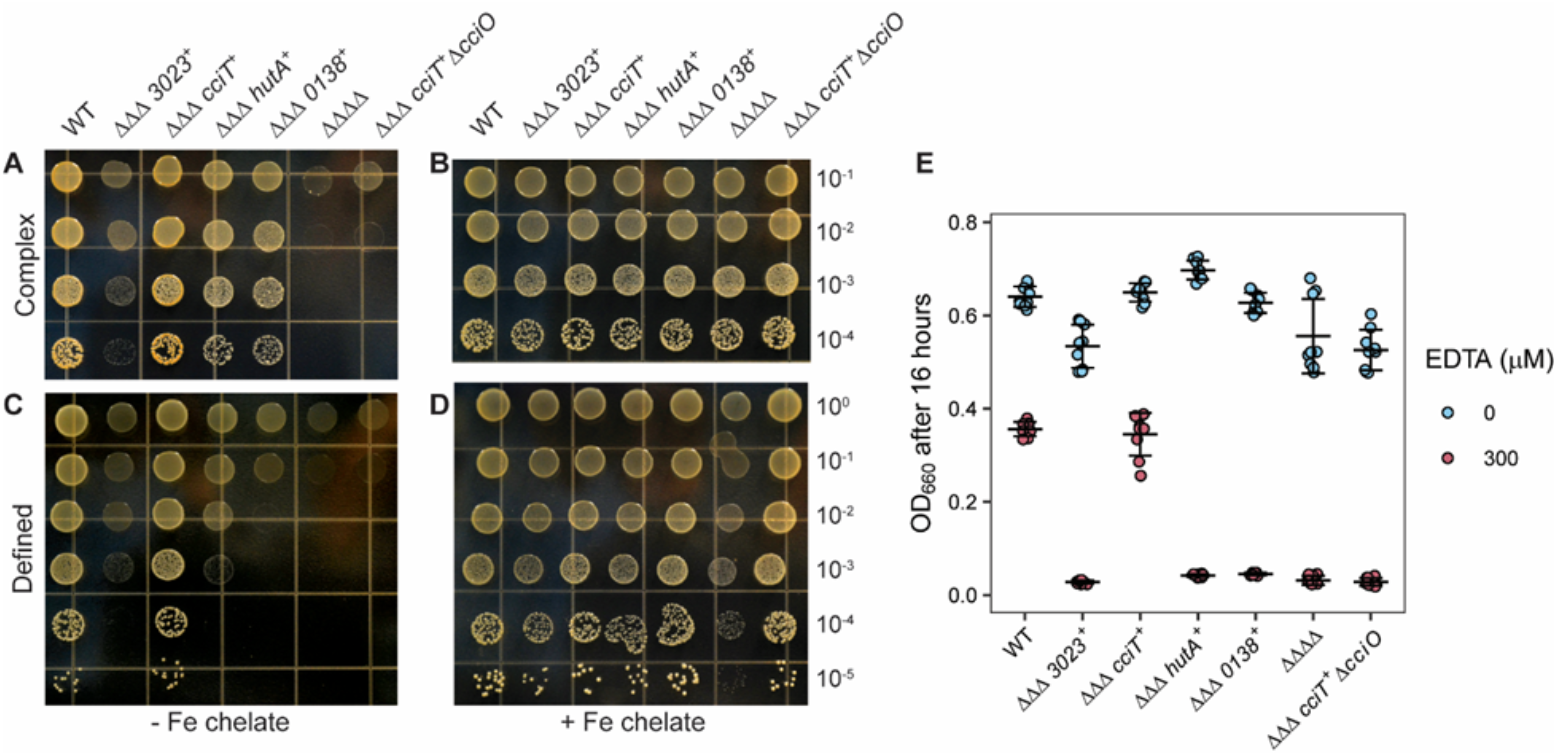
*cciT* alone is sufficient to support wild type-like growth on solid medium; this function requires *cciO*. (WT), strains lacking three of the four Fur-regulated TBDT genes (ΔΔΔ CCNA_03023^+^, ΔΔΔ cciT^+^, ΔΔΔ hutA^+^, and ΔΔΔ CCNA_00138^+^), and a strain lacking all four Fur-regulated TBDT genes (ΔΔΔΔ) were serially diluted 10-fold and spotted onto solidified complex PYE medium (**A, B**) or defined M2X medium (**C, D**), without (**A, C**) or with 10 μM Fe•EDTA (**B, D**). (**E**) Optical density at 660 nm of all strains after growth for 16 h in complex liquid PYE medium without (blue) or with 300 μM EDTA (pink). The mean ± standard deviation of three independent trials of three independent cultures (n = 9), are overlayed with the individual data points.

### CciT transporter function requires the CciO dioxygenase

CciO is commonly encoded from a chromosomal locus adjacent to *cciT* in Proteobacteria (Figure 2) and has a functional role in the maintenance of cellular iron levels (Figures 4–6). Given the similar growth phenotypes of strains lacking either *cciT* or *cciO* (Figures 3, 5, 6 & S2), we postulated that the CciO dioxygenase was specifically required for CciT-supported growth. To assess the functional relationship between *cciO* and *cciT*, we deleted *cciO* from the triple TBDT deletion strain, retaining *cciT* as the only intact TBDT gene (designated ΔΔΔ *cciT*^+^ Δ*cciO*). The growth defect of this strain was comparable to that of the ΔΔΔΔ strain lacking all four Fur-regulated TBDT genes when grown on complex solid medium, defined solid medium, and in liquid defined medium (Figures 6 and S2D). Thus, the ability of CciT to support *Caulobacter* growth requires CciO. To determine if the function of CciO is specific to the CciT transporter or if it plays a broader role in iron acquisition, we deleted *cciO* in other mutant strains harboring only one of the four Fur-regulated TBDT genes. We thus generated the strains ΔΔΔ *138*^+^ Δ*cciO*, ΔΔΔ *3023*^+^ Δ*cciO*, and ΔΔΔ *hutA*^+^ Δ*cciO*. The growth advantage provided by each of these single transporter genes was unaffected by deletion of *cciO* (Figure S3C). We conclude that there is a specific iron transport relationship between the dioxygenase CciO and the TonB-dependent transporter CciT.

### CciT and CciO support iron acquisition under chelation stress

CciT and CciO are EDTA resistance factors that function together as a major *C. crescentus* iron-acquisition system. However, the mechanism by which EDTA treatment compromised growth of the *cciT* and *cciO* mutants remained unclear. To address this question, we first queried the specific contribution of each Fur-regulated TBDT gene to EDTA resistance by measuring growth in complex (PYE) liquid medium containing 300 µM EDTA. CciT fully and uniquely supported *C. crescentus* growth at this EDTA concentration, and its ability to execute this function required *cciO* (Figure 6E). To further assess iron-acquisition ability of the WT, Δ*cciO*, and Δ*cciT* strains in medium with 300 µM EDTA, we supplemented liquid medium with increasing concentrations of FeCl_3_ (1, 10, 50, and 100 µM) and measured growth. Supplementation with FeCl_3_ enhanced the growth of all strains in a dose-dependent manner (Figure 7A), but to different degrees. The addition of as little as 10 µM FeCl_3_ was sufficient to maximize the growth enhancement of WT cultures, whereas the growth of the Δ*cciO* and Δ*cciT* cultures was only modestly enhanced at this concentration. The Δ*cciO* and Δ*cciT* mutant strains required five times more ferric iron (50 µM FeCl_3_) than WT to achieve near maximal growth enhancement. We conclude that the addition of 300 µM EDTA to the medium impairs the ability of *cciT* and *cciO* mutants to acquire ferric iron.

**Figure 7.**
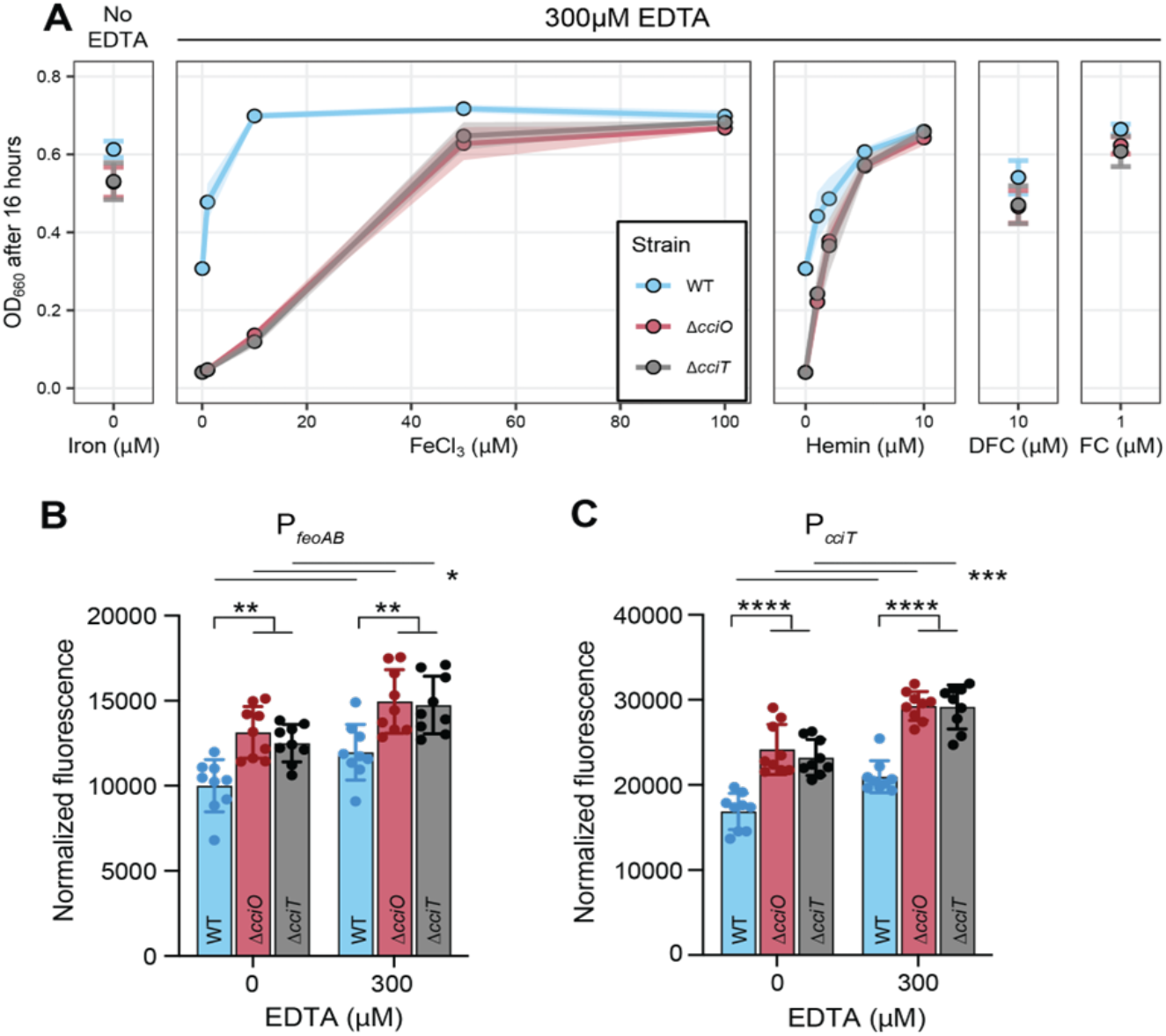
*ΔcciO* and *ΔcciT* have diminished ferric iron–acquisition ability in the presence of EDTA. **(A)** Optical density at 660 nm of wildtype (WT), ΔcciO, and ΔcciT strains after 16 h of growth in PYE medium without (far left) or with 300 µM EDTA (center and right) with increasing concentrations of FeCl_3_ or hemin, 10 mM desferrichrome (DFC), or 1 mM ferrichrome (FC) as indicated on the X-axis. Values represent mean ± SD (as shaded area) of at least six independent replicates. **(B&C)** Fluorescence output (normalized to cell density) of WT, ΔcciO, and ΔcciT strains carrying an mNeonGreen reporter cassette under the control of the Fur-regulated feoAB (**B**) or cciT promoters (**C**). Cells were grown for 1.5 hours in liquid PYE medium without or with 300 µM EDTA. Values are means ± SD of 9 independent cultures assayed over three different days.Iindividual data points are also shown. A two-way ANOVA followed by Tukey’s post-hoc test was used to assess differences between strains (lower brackets) and between treatments (upper lines). * p < 0.05; ** p < 0.01, *** p < 0.001, **** p < 0.0001).

EDTA treatment was reported to lower the level of cellular iron, manganese, and zinc in *E. coli* (16). Transcriptional reporter assays showed that EDTA treatment increases transcription of Fur-regulated genes in the WT *C. crescentus* strain (Figure 7B & C), and that EDTA-induced iron limitation is more severe in Δ*cciO* and Δ*cciT* mutants, as we detected higher normalized fluorescence from the reporters in the mutant backgrounds than in WT (Figures 1C, 7B & C). While *C. crescentus* can use Fe•EDTA as an iron source, we propose that CciT–CciO enables transport of an undefined Fe^III^-siderophore complex at a lower concentration than iron from Fe•EDTA. From this model, we predicted that adding a chelated iron source that is transported through an alternative high-affinity transport system would rescue the growth defects of strains lacking the CciO–CciT system when EDTA is present. Indeed, adding 1 µM of hemin—an iron-porphyrin complex transported by HutA (67)—partially restored the growth of the Δ*cciO* and Δ*cciT* mutant strains cultivated in a complex medium with 300 µM EDTA (Figure 7A), while 10 µM hemin restored growth to WT levels. We further tested this model using the hydroxamate siderophore desferrichrome (DFC), which is efficiently transported by *C. crescentus* at sub-nanomolar concentrations when bound to iron (67). DFC has a higher overall stability constant with Fe^III^ than EDTA (68) and can thus compete with EDTA for available ferric iron in the medium. Adding 10 µM desferrichrome to a complex liquid medium containing 300 µM EDTA restored the growth of the mutant strains to near WT levels, while adding 1 µM of DFC pre-loaded with Fe^III^ (i.e. ferrichrome, FC) fully rescued the growth of the Δ*cciO* and Δ*cciT* mutants challenged with 300 µM EDTA (Figure 7A). We conclude that *cciT* and *cciO* enable the uptake of iron under EDTA chelation stress. However, the growth of Δ*cciT* and Δ*cciO* mutants in the presence of EDTA can be rescued by chelated iron transport through alternative high-affinity systems, such as those for ferrichrome and hemin.

### Assessing the role of the CciT–CciO system in the import of catechol siderophores

A phylogenetic analysis of *C. crescentus* and *E. coli* TBDTs revealed a close relationship between CciT and Fiu (67), which transports monomeric catechol-based siderophores in *E. coli* (69). As presented above, Fiu is encoded from a locus adjacent to a 2OG/Fe^II^-dependent dioxygenase gene homologous to *cciO* and functions as the outer membrane transporter for the catechol-cephalosporin antibiotic cefiderocol (67). A function for *C. crescentus* CciT as a Fiu/PiuA-like catechol transporter would imply that deletion of *cciT* would result in cefiderocol resistance, as reported in resistant clinical isolates of several bacterial pathogens (35, 70). Unexpectedly, WT *C. crescentus* was highly resistant to cefiderocol treatment, and deletion of either *cciT* or *cciO* increased this sensitivity (Figure S6). Supplementing the medium with 10 µM Fe•EDTA attenuated the cefiderocol sensitivity of the mutants. We conclude that CciT does not transport cefiderocol and propose that another transport system, upregulated upon iron limitation in the Δ*cciO* and Δ*cciT* mutant backgrounds, transports this catechol-containing antibiotic into *Caulobacter* cells. This unknown cefiderocol/catechol transport system is apparently not one of the other three Fur-regulated TBDTs, as the strains harboring a single Fur-regulated TBDT gene and the quadruple deletion strain (Δ Δ Δ Δ) were all sensitive to cefiderocol (Figure S7).

We tested other known catecholate siderophores as possible CciT substrate(s), including 2,3′-dihydroxybenzoic acid (2,3-DHBA) (69), chlorogenic acid (66), and protocatechuate (3,4-DHBA), which is known to be catabolized by *C. crescentus* (72). In a defined liquid medium, the addition of Fe^III^ complexed with any of these three catecholates at a 1 µM Fe^III^:10 µM catecholate ratio enhanced growth of all multi-TBDT gene deletion strains relative to 1 µM Fe^III^ provided as FeCl_3_, although iron complexed to 2,3-DHBA or chlorogenic acid enhanced growth more than protocatechuate (Figure S8). The growth phenotype of ΔΔΔ*cciT*^*+*^*ΔcciO* matched that of the ΔΔΔΔ strain in most conditions, consistent with a model in which CciT function requires CciO. The one exception was in medium containing Fe^III^ mixed with 2,3-DHBA, where catechol-enhanced growth did not require *cciO* (Figure S8).

The cefiderocol resistance and Fe^III^-catecholate growth supplementation data provided evidence that *C. crescentus* can transport catecholates through multiple routes at the tested concentrations. A specific growth enhancement of a particular strain (or strains) carrying single Fur-regulated TBDT genes was not evident for any of the tested catecholates (Figure S8B), and the data clearly show that the *cciT*–*cciO* locus conferred the greatest growth advantage under conditions with no added iron. This result indicates that the CciT–CciO system either supports acquisition of some form of chelated iron present in M2 defined medium or that CciT transports a siderophore produced by *C. crescentus* itself, a possibility that we discuss below.

### Growth and EDTA sensitivity of Δ*cciO* and Δ*cciT* in a natural environment context

The *cciT*–*cciO* locus encodes a major iron-acquisition system in *C. crescentus* that supports fitness and EDTA resistance in traditional laboratory media. However, the natural habitats of *Caulobacter* spp. are primarily aquatic and soil ecosystems (19). As such, we assessed the consequences of *cciT*–*cciO* gene deletion and EDTA treatment on *C. crescentus* growth in *bona fide* freshwater collected from two lakes in Michigan, USA: a developed recreational lake (Lake Lansing) and a Laurentian Great Lake (Lake Huron). When studying bacterial growth in a complex natural medium, the impact of microbially-derived metabolites, such as siderophores, should be considered. Indeed, we have shown that desferrichrome, a common environmental siderophore (73) with an Fe^III^ stability constant that is higher than EDTA, alleviates growth inhibition caused by EDTA treatment (Figure 7A). Given the likely presence of siderophores in our freshwater samples (74, 75) and the extensive repertoire of TBDTs encoded by *C. crescentus*, we hypothesized that *C. crescentus* growth in lake water would be less dependent on *cciT*-*cciO* and less impacted by the presence of EDTA. To test this, we cultivated the WT, Δ*cciO*, and Δ*cciT* strains in water from Lake Lansing or Lake Huron, both with and without 300 µM EDTA, and measured strain growth by enumerating CFUs.

The populations of *C. crescentus* cells grown in these natural water samples underwent ∼6–8 doublings, reaching a higher density in water collected from Lake Lansing compared to that from Lake Huron (Figure 8). Phosphorus levels were similar between lake samples (Figure S10 and Table S4), but total organic carbon (7.7 versus 4.5 mg/mL) and Kjeldahl nitrogen levels (0.80 versus 0.33 mg/mL) were higher in water samples from Lake Lansing (Table S5), which likely explains higher levels of *C. crescentus* growth in those water samples. As predicted, we observed no significant differences in growth between WT cells and the *ΔcciO* or *ΔcciT* mutant strains in these lake water samples regardless of addition of 300 µM EDTA (Figure 8). 1000 µM EDTA attenuated growth of the WT strain in Lake Lansing and Lake Huron water while 10,000 µM EDTA killed all cells (Figure S9). These results demonstrate that *C. crescentus* (a) is less sensitive to EDTA chelation stress in *bona fide* fresh-water conditions and (b) can use other iron sources in these freshwaters beyond the form of iron that is imported by the CciT–CciO system.

**Figure 8.**
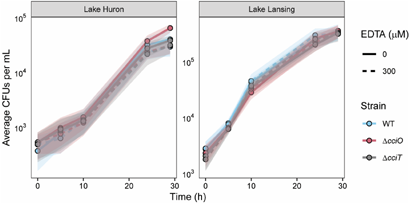
Growth of mutant strains in water collected from Lake Huron and Lake Lansing is not affected by the addition of EDTA. WT, *ΔcciO*, and *ΔcciT* strains inoculated into water collected from Lake Huron (left) or Lake Lansing (right) that was filtered before inoculation, with (dashed lines) and without (solid lines) 300 µM EDTA added. CFUs were enumerated after 0, 5, 10, 24, and 29 h of growth. Values represent means ± SD (as shaded areas) of three replicate cultures generated on the same day.

### Quantitative elemental analysis of growth media and the relationship between cation profile and EDTA toxicity

We further considered how differences in the chemistry of laboratory media and lake water samples may contribute to the differential growth and EDTA sensitivity of *C. crescentus* across these environments, and therefore determined the elemental profiles of non-supplemented liquid M2X defined medium, liquid PYE complex medium, and water collected from the two lakes by ICP-MS-QQQ. M2X contained the expected concentrations of the major medium components (Figure S10 and Table S4). Sodium, potassium, and phosphorus levels were ∼2–10 times lower in PYE medium relative to M2X, although the levels in PYE were still

∼3–30 times higher than in lake water (Table S4 and Figure S10). Sulfur levels were the highest in PYE medium (∼2 mM) among all tested media and were substantially lower in the lake water samples. A difference in sulfur content between Lake Lansing and Lake Huron was reflected in the higher level of measured sulfate ions in Lake Huron water samples (16 mg/L) relative to Lake Lansing (<5 mg/L). The contents of magnesium, copper, nickel, zinc, cobalt, chromium, selenium, and iron were all higher in PYE medium than in M2X medium and lake water. The level of iron in non-supplemented M2X defined medium approximated that measured in the water samples from Lake Lansing (80 nM) and Lake Huron (187 nM), while the concentration of iron in PYE medium was significantly higher (635 nM). Calcium concentrations in the lake waters were ∼700 µM, which is congruent with that in PYE medium and ∼2 times higher than in M2X medium. The toxic metals chromium and cadmium were more abundant in laboratory media than in the lake waters (Table S4 and Figure S10). Cadmium levels were substantially higher (by a factor of 16) in M2X defined medium than in complex PYE medium and were below the instrument detection limit in the lakes. We are uncertain what component(s) of M2 base medium contain higher levels of cadmium.

EDTA is a broad-spectrum metal chelator that associates with many different cations. Thus, the particular cation profile of a medium will influence the effect of EDTA on bacterial growth. We examined whether supplementing select cations could counteract the detrimental effects of 300 µM EDTA on *C. crescentus* growth in liquid PYE complex medium. Adding 2 mM Ca^II^ or 1 µM Fe^III^ partially rescued growth in the presence of EDTA. Similarly, the addition of 300 µM Zn^II^, Mn^II^, or Ni^II^ mitigated the growth-limiting effect of EDTA, although the addition of Zn^II^ or Ni^II^ alone (without EDTA) was toxic to *Caulobacter* (Figure S11). Altogether, our data provide evidence that the spectrum of cations along with the availability of chelated forms of iron (e.g. ferrichrome, heme, or other Fe-siderophores) will influence EDTA toxicity in freshwater.

## Discussion

A forward genetic screen for *C. crescentus* genes that affect fitness in the presence of the broad-spectrum synthetic chelator EDTA uncovered a TonB-dependent transporter (CciT) and a 2OG/Fe^II^-dependent dioxygenase (CciO) that coordinately function to support cellular iron homeostasis. Our study defines *cciT–cciO* as encoding key iron-acquisition proteins and demonstrates the importance of these coconserved, Fur-regulated genes in sustaining microbial growth under iron limitation and chelation stress.

*Caulobacter* has multiple routes through which it can acquire iron (67), but the CciT–CciO system provides the largest fitness advantage of all the Fur-regulated iron-acquisition systems in both complex and defined media. Deletion of any other Fur-regulated TBDT gene had no effect on growth (Figure 3), and no other single Fur-regulated TBDT besides CciT was sufficient to support WT-like growth on laboratory media (Figures 6 and S2D). In a complex medium containing EDTA, CciT uniquely supported fitness among the Fur-regulated TBDTs. The function of CciT in these treatment conditions strictly required the presence of the dioxygenase gene *cciO* (Figure 6E).

### *Transcriptional signatures of iron limitation in* ΔcciT *and* ΔcciO

Deletion of either gene in the *cciT-cciO* operon led to increased transcription of a small set of genes, most of which are repressed by Fur, including *cciT* and *cciO* themselves (Figure 4 and Table S3). Several of the regulated genes were previously shown to be activated in response to treatment with the iron chelator 2,2’-dipyridyl (DP) (62, 67), though the transcriptional response to *cciT* or *cciO* deletion was less pronounced than the acute iron starvation response induced by DP. These transcriptomic data support the conclusion that loss of *cciT* or *cciO* function leads to reduced intracellular iron levels in both complex and defined media.

Notably, several uncharacterized genes that were modestly but significantly upregulated in Δ*cciT* and Δ*cciO* strains compared to WT have been linked to iron acquisition in other genera. For instance, genes from a predicted operon encoding a flavin-dependent reductase, a PepSY-family membrane protein (COG3295), and DUF2271 and DUF4198 domain proteins (locus CCNA_03155-03159) were significantly upregulated. Transposon insertions in these genes, which are proposed to facilitate periplasmic ferric iron reduction (47), decreased *C. crescentus* fitness in the presence of EDTA (Figure 1 and Table S1). Experimental data from *Vibrio cholerae* further support a role for these COG/DUF proteins in iron acquisition, with COG3295 protein VciB promoting Fe^II^ transport by facilitating Fe^III^ reduction (48, 49). Additionally, transcriptome data from *Alteromonas macleodii* suggest that DUF2271 and DUF4198 proteins are also involved in iron acquisition (50).

### Quantitative elemental analysis of the environment

In an effort to understand the variable sensitivity of *C. crescentus* to loss of *cciT* or *cciO* function in different media environments, we determined the elemental profiles of the artificial and natural media in which strains were cultivated (Table S4). To our knowledge, this study provides the first quantitative inorganic analysis of *Caulobacter* cultivation media. However, it is important to note that these results are influenced by the source of the media supplies, as the levels of trace elements can vary between different manufacturers and lots. These data, when viewed in the context of accompanying analytical data from two Michigan lakes that supported the growth of *C. crescentus* (Figure 8), offer an opportunity to reconsider effects of the growth media that are used to evaluate the physiology and ecology of this important environmental bacterium. While the balance of available carbon, nitrogen, and phosphorus was orders of magnitude higher in defined (M2X) and complex (PYE) media than in lake waters (Figure S10 and Table S5), the ensemble of inorganic elements varied considerably. Sodium, potassium, sulfur, and nickel levels were 1–2 orders of magnitude higher in laboratory media than in waters of Lake Lansing or Lake Huron. The concentrations of essential metals including iron, magnesium, copper, cobalt, and zinc were comparable between lake water samples and M2X defined medium (without added iron supplements) but were 3–300 times higher in complex medium (Table S4 and Figure S10). Measured levels of calcium, manganese and molybdenum in the lakes were better reflected in complex medium than in defined medium. The levels of the toxic metals cadmium and chromium were both significantly higher in laboratory media than in the lake waters, which may influence *Caulobacter* physiology under certain experimental conditions. As the field continues to explore *Caulobacter* nutrient acquisition and stress physiology in environmental contexts, it may be useful to formulate media that have an inorganic composition that is more aligned with the composition of lake waters from this study.

### Interactions of EDTA with Caulobacter and its growth medium

Experiments assessing the interactions between *C. crescentus*, EDTA, and its growth media have highlighted the redundancy and versatility of its iron-acquisition strategies. *C. crescentus* can use Fe•EDTA as an iron source (73), but we propose that *C. crescentus* does not utilize Fe•EDTA as efficiently, or at as low a concentration, as it does the *bona fide* substrate of CciT. In support of this model, we showed that CciT and CciO are strictly required for growth in a complex medium containing a concentration of EDTA that only partially limits growth of WT cells (Figure 6). We further demonstrated that *C. crescentus* CciT and CciO enable iron acquisition at low ferric iron concentrations, even when EDTA was present in high molar excess (Figure 7). Interactions between EDTA, the culture medium, and *Caulobacter* cells are complex and dependent on the chemical composition of the medium (Figure 7, S10 & S11). Our data suggest redundancy in the ability of the multiple *Caulobacter* iron-acquisition systems to use not only Fe•EDTA but also a diverse set of Fe^III^-catechol substrates when these compounds are available at low micromolar concentrations (Figures S6– S8). The CciT–CciO system is apparently the major iron import system in complex and defined laboratory media lacking chelated iron, but the WT-like growth phenotypes of the Δ*cciT* and Δ*cciO* mutants in lake water (with or without 300 μM EDTA) (Figure 8) reflect the adaptability of this common freshwater and soil microbe in acquiring essential nutrients in chemically complex natural habitats. The mechanism by which Lake Lansing and Lake Huron waters (effectively) buffer *Caulobacter* against EDTA treatment is not known, but there are almost certainly multiple chemical factors involved. The distinct metal ion profiles (Table S4 & Figure S10) and the highly complex array of inorganic and organic iron- and other metal-binding ligands present in natural waters (74–77), likely mitigate the biological effect of EDTA.

### On the CciT/CciO substrate(s)

The outer membrane transporter CciT and the cytoplasmic dioxygenase CciO coordinately support robust growth of *C. crescentus* on both solid and liquid defined media (Figures 6 and S2). Given the strongly reduced growth of Δ*cciT* and Δ*cciO* relative to the WT strain in mineral defined media supplemented with FeCl_3_, we propose that CciT imports ferric iron bound to a substrate that is not present in the medium and is produced by *C. crescentus* itself. The identity of this postulated compound is not known, but the evolutionary relationship between CciT, *E. coli* Fiu (67), and *Pseudomonas* spp. PiuA suggests that it would have a catecholate-like structure (Figure 9). Efforts to identify siderophore production by *C. crescentus* using colorimetric (78) and other analytical approaches have been unsuccessful to date, and the catechol supplementation and cefiderocol resistance data presented herein (Figures S6–S8) support the existence of multiple routes for catecholate entry at the tested concentrations.

**Figure 9:**
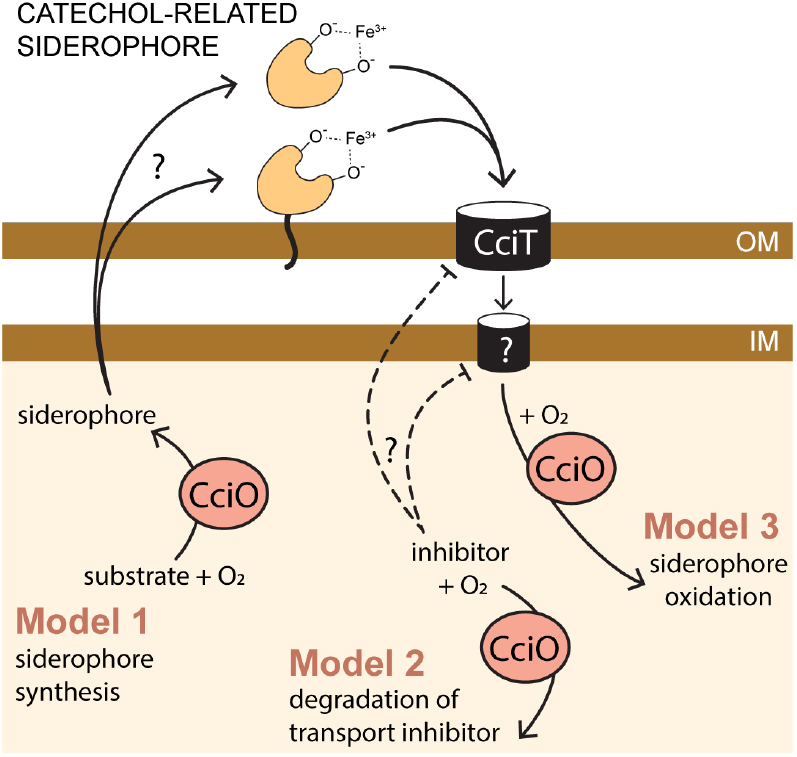
CciT and CciO function together to support iron homeostasis. *cciT* is a Fur-regulated gene encoding an outer membrane transporter related to known Fe^III^•catechol transporters. Iron acquisition via CciT specifically requires CciO, a conserved dioxygenase. CciO may be required to synthesize the substrate of CciT (Model 1), degrade an inhibitor of CciT-dependent transport (Model 2), or oxidize the substrate of CciT (Model 3). The postulated iron-binding substrate of CciT may be cell-associated, as reported in many marine bacteria (89). The inner membrane transporter is not known.

The growth (Figures 3 & 6) and intracellular iron limitation phenotypes (Figure 5) of the Δ*cciO* strain matches those seen in Δ*cciT*, which provides evidence that the iron import function of CciT at the outer membrane requires the intracellular dioxygenase activity of CciO. Proteomic analysis of fractionated membranes demonstrated that CciT is produced and properly targeted to the outer membrane in the absence of CciO (Figure S4). Thus, a model in which CciT translation and/or targeting requires CciO activity can be ruled out. It is possible that CciO is involved in the biosynthesis of a siderophore, although we would not have expected to identify *cciO* mutants in our pooled RB-TnSeq screen (Figure 1) if CciO were required to synthesize a “shared good” that was secreted to the medium. If CciO is involved in the biosynthesis of a Fe^III^-binding siderophore that is tethered to the outside of the cell as observed in some marine species (79) (Figure 9), then we may expect decreased fitness of *cciO* mutants in a pooled mutant screen, as we observed. A membrane-associated siderophore model is also consistent with ICP-QQQ-MS data showing reduced total cellular iron in Δ*cciO* but not in Δ*cciT* (Figure 4D) Homologs of CciO in *P. aeruginosa* and *E. coli* (known as PiuC or YbiX) have been suggested to transform the substrate transported by their respective adjacently encoded TBDT (80). CciO may participate in the catabolism of this postulated iron-bound siderophore, which could release the bound iron or help to sustain a favorable siderophore concentration gradient for import. Alternatively, CciT and CciO may not have the same (or even related) substrates. This study demonstrates a clear functional link between this widely co-conserved gene pair (Figure 2), and it is our hope that future studies will define the molecular connections between outer membrane iron transport and cytoplasmic diox-ygenase activity.

## Acknowledgements

We thank Bob Hausinger and Howard Shuman for helpful comments on this manuscript, and Claire Veille, Michaela TerAvest, and Bob Hausinger for insight and guidance over the course of this study. Research reported in this publication was supported in part by the National Institute of General Medical Science of the National Institutes of Health under award numbers R35GM131762 to S.C., P41GM135018 and R01GM038784 to T.V.O., and by Army Research Office contract W911NF2210105 to S.C.

## Materials and Methods

### Growth conditions and strain construction

*C. crescentus* strains were grown at 30°C in one of several types of media as indicated. The peptone-yeast extract (PYE) medium (0.2% [w/v] peptone (Fisher Bioreagents, Lot No. 225155), 0.1% [w/v] yeast extract (Fisher Bioreagents, Lot No. 220635), 1 mM MgSO_4_ (Fisher Chemical, Lot No. 183674), 0.5 mM CaCl_2_ (Fisher Chemical, Lot No. 117031)) is best described as a complex medium. In contrast M2X, is a defined medium based on M2 salt medium (6.1 mM Na_2_HPO_4_ (Aldrich Chemical Company, Inc., Lot No. 12030MN), 3.9 mM KH_2_PO_4_ (Fisher Bioreagents, Lot No. 141122), 9.3 mM NH_4_Cl (Fisher Chemical, Lot No. 156427), 0.25 mM CaCl_2_, 0.5 mM MgSO_4_, 10 µM Fe•EDTA chelate (Sigma Life Science, Lot No. RNBD1641)) that has 0.15% (w/v) xylose (Acros Organics, Lot No. A0408426) added as a carbon source. The Fe•EDTA chelate in M2X, and occasionally used to supplement PYE, is a 1:1 molar mix of FeSO_4_ and EDTA (ferrous sulfate chelate solution kept at 4°C, Sigma-Aldrich, F0518). However, the iron source in M2X was modified in some experiments, or omitted, as indicated. When ferric iron was used, a 100 mM FeCl_3_ aqueous stock (pH ∼3.0) kept at ™20°C was thawed and used to make a working 10 mM stock that was added just before use to the final concentrations indicated. A 10 mM hemin stock solution was prepared in 1 M NaOH and diluted to the indicated final working concentrations. When Fe•EDTA was added to solid medium, it was added after autoclaving. When an iron source (Fe•EDTA, FeCl_3_, hemin, desferrichrome (*Ustilago sphaerogena*; Sigma-Aldrich)) was added to liquid media, it was added to sterile media just before bacterial inoculation. The water used for the preparation of all media was ultra-purified using a Barnsted GenPure water purification system (ThermoFisher). *E. coli* strains were grown at 37°C in LB medium (1% [w/v] tryptone, 0.5% [w/v] yeast extract, 1% [w/v] NaCl). Growth media were solidified by the addition of 1.5% (w/v) agar when necessary. Antibiotics were used at the following concentrations in liquid and solid media as appropriate: *C. crescentus*, 5 µg/mL or 25 µg/mL kanamycin, 1 µg/mL or 2 µg/mL chloramphenicol, 1 µg/mL or 2 µg/ml tetracycline, 20 µg/mL nalidixic acid; *E. coli*, 50 µg/mL kanamycin, 20 µg/mL chloramphenicol, 10 µg/mL tetracycline.

Cells were also grown in environmental water samples from Lake Lansing and Northern Lake Huron (Hammond Bay). These samples were collected in high-density polyethylene analysis bottles provided by the Michigan Department of Environment, Great Lakes, and Energy (EGLE). Bottles were pre-rinsed with lake water from the sample site before sampling. Lake Huron waters were collected on November 10, 2023, in Presque Isle County, Michigan, USA, at 45°31’01.9”N 84°07’09.8”W. Lake Lansing waters were collected on February 14, 2024, from Ingham County, Michigan, USA, at 42°45’19.2”N 84°24’17.2”W. A quantitative elemental profile of these environmental samples was determined as described below.

Standard molecular biology techniques were used to construct all plasmids. Detailed information on strains, plasmids, and primers can be found in Table S6. To create plasmids for in-frame deletion allele replacements, the regions flanking the target gene were cloned into pNPTS138. For the generation of fluorescent transcriptional reporter plasmids, promoter regions (400–500 bp upstream of the open reading frame) were cloned into into the vector pPTM056 (81). For complementation, the entire coding sequences with their respective promoter regions were cloned, without generating a fluorescent fusion, into pMT603 (pXGFPC5) (21), which integrates at the *xylX* locus.

All plasmids were individually introduced into the *C. crescentus* CB15 strain either by electroporation or by tri-parental mating. For allele replacements, a double recombination strategy was used to select merodiploid strains harboring each relevant pNPTS-based plasmid with kanamycin resistance, followed by *sacB* counter selection on PYE plates containing 3% (w/v) sucrose. PCR was used to evaluate colonies that were sucrose-resistant and kanamycin-sensitive to identify those colonies harboring the null allele. For the construction of strains with multiple gene deletions for TonB-dependent transporter genes, counterselection was performed on PYE sucrose plates supplemented with 10 µM Fe•EDTA (Sigma-Aldrich, F0518).

### Tn-himar-seq to assess gene contributions to fitness in the presence of EDTA

An aliquot of a *C. crescentus* CB15 barcoded Tn-himar mutant library previously reported (39, 40) was grown in 5 mL of liquid PYE medium containing 5 µg/mL kanamycin for 8 h. One milliliter of this pre-culture was set aside as a reference, and four replicate cultures of 2 mL each in liquid PYE medium, alone or with 300 µM EDTA, were inoculated to a starting optical density at 660 nm (OD_660_) of 0.01. The cultures were incubated in 100 mm ’ 14 mm glass culture tubes with shaking at 200 RPM at 30°C for 17 h, and 1 mL from each culture was collected by centrifugation for 3 min at 16,000*g*. The supernatants were discarded, and the cell pellets were resuspended in 10–20 µL of water and frozen at ™20°C.

Barcode abundances were determined following the method developed by Wetmore et al (29). Briefly, barcodes were amplified using Q5 polymerase (New England Biolabs) in a 20-µL reaction containing 1’ Q5 reaction buffer, 1’ GC enhancer, 0.8 units Q5 polymerase, 0.2 mM dNTPs, 0.5 µM of each primer, and 1 µL of the cell suspension, using the following amplification conditions: 98°C for 4 min, 25 cycles of (98°C for 30 sec, 55°C for 30 sec, and 72°C for 30 sec), 72°C for 5 min, followed by a hold at 4°C. The primer set consisted of a universal forward primer, Barseq_P1, and a uniquely indexed reverse primer, Barseq_P2_ITxxx, with the index number denoted by xxx (see ref (29)). The barcode amplification products were pooled and sequenced as 50-bp single-end reads on an Illumina MiSeq instrument, using Illumina TruSeq primers. Barcode sequence analysis was carried out using the fitness calculation protocol of Wetmore et al (29). Barcodes from each sample were counted and compiled using MultiCodes.pl and combineBarSeq.pl scripts (29). A barcode table was generated, from which the FEBA.R script was used to calculate fitness score relative to the reference starter culture.

### Pangenome analysis

The genomes of 69 *Caulobacter* strains were analyzed using Anvi’o version 7.1 (60). The genomes incorporated into the analysis, as detailed in Table S2, were downloaded from the NCBI RefSeq database. These genomes were reformatted with the ‘anvi-scriptreformat-fasta’ tool and integrated into an Anvi’o contigs database via ‘anvi-gen-contigs-database’. Open reading frames (ORFs) were predicted using Prodigal (86). Predicted genes received functional annotations based on COG terms (84), and these annotations were subsequently appended to the contigs database. Gene clusters were delineated with the mcl algorithm (85), and a pangenome was constructed using ‘anvi-pan-genome’ with the parameters --mcl-inflation set to 6 and --min-occurrence set to 10.

### Neighborhood analysis

The sequence of the CciO (encoded by gene locus CCNA_00027) protein was analyzed using the PSI-BLAST tool, set to default parameters, against the NCBI RefSeq database while excluding sequences from uncultured/environmental samples. Proteins that had greater than 95% query coverage and more than 65% identity were selected. The accession IDs of these proteins were submitted to the WebFLaGs server for neighborhood analysis, available at http://www.webflags.se/ (61). For brevity of presentation, a manually curated set of diverse loci is presented in Figure 2.

### Serial dilution plating experiments

Overnight cultures grown in M2X medium with 10 µM FeSO_4_•EDTA were centrifuged at 16,000*g* for 3 min at room temperature. Cell pellets were resuspended in iron-free liquid M2X medium to an OD_660_ of 0.2 and serially diluted 10-fold. Using a multi-channel pipette, 5 µL from each dilution was spotted onto an agar plate. After the droplets were allowed to dry, the plates were incubated at 30°C for 2–3 days.

### Growth measurements

A single colony from a strain grown on a PYE plate was inoculated into 2 mL of liquid PYE medium and grown overnight. The next day, the cultures were diluted to an OD_660_ of 0.15 and grown for 2 h before being used to inoculate fresh cultures for the desired experimental condition (medium) at an OD_660nm_ of 0.01. Cultures were then incubated at 30°C with shaking at 200 rpm. After 16 h of growth, the optical density of the cultures was measured.

### Measurement of growth in liquid M2X medium

Primary cultures in liquid M2X medium were inoculated from cells freshly grown on solid plates to an initial OD_660_ of 0.02–0.10 and grown overnight. These overnight cultures were diluted to an OD_660_ of 0.002 and allowed to grow for another day. The following morning, cells were collected by centrifugation and resuspended in iron-free liquid M2X medium. These cell suspensions were used to inoculate into various media at a starting OD_660_ of 0.01. Cultures were grown at 30°C with shaking at 200 rpm. Growth was periodically monitored by measuring OD_660_. In conditions where iron was bound to a siderophore, FeCl_3_ was combined with chelator in a 1:10 molar ratio prior to being added to the medium (see catecholate supplementation section below).

### Cell size analysis

Wild-type and mutant *C. crescentus* cells were grown in PYE broth overnight. Starter cultures were used to inoculate 12 mL of PYE broth in 20 mL glass tubes at an OD_660_ of 0.01. Cultures were grown for 6 hours at 30°C shaking at 200 RPM. To sample the cultures, 1 µl of cells were spotted on an agarose pad on a cover slide and imaged on a Leica DMI 6000 microscope in phase contrast with an HC PL APO 63x/1.4 numeric aperture (NA) oil Ph3 CS2 objective. Images were captured with an Orca-ER digital camera (Hamamatsu) controlled by Leica Application Suite X (Leica). Cell areas were measured from phase contrast images using BacStalk (86).

### Transcriptome sequencing (RNA-seq)

The WT strain CB15 and the Δ*cciO* (lacking CCNA_00027) and Δ*cciT* (lacking CCNA_00028) mutant strains were grown overnight at 30 C in 2 mL of liquid M2X medium containing the standard 10 µM Fe•EDTA chelate concentration. These cultures were used to inoculate 10 mL of liquid M2X medium to a starting OD_660_ of 0.001 and then incubated overnight. After the OD_660_ exceeded 0.1, the cells were collected by centrifugation, rinsed with iron-free liquid M2X medium, and resuspended in 10 mL of liquid M2X medium containing 10 µM FeCl_3_. The cell suspension was adjusted to an OD_660_ of 0.1 before being incubated at 30°C with shaking at 200 RPM for 2.5 h. Cells were harvested by centrifugation and stored at ™80°C until RNA extraction. RNA was extracted using TRIzol (Invitrogen) as previously described (87). RNA-seq libraries were prepared utilizing an Illumina TruSeq Stranded RNA Kit and sequenced on an Illumina NextSeq 500 instrument at SeqCoast Genomics (Portsmouth, NH). The sequencing data were processed with the CLC Genomics Workbench 20 (Qiagen), aligning reads to the *C. crescentus* NA1000 genome (Genbank accession CP001340). RNA sequencing data are available in the NCBI GEO database under accession number GSE274268.

### Analysis of transcription using fluorescent reporters

Strains harboring fluorescent transcriptional reporter plasmids to monitor the activity of the *feoB* or *cciT* promoters (fused to mNeon-Green) were cultured overnight in triplicate at 30°C in liquid PYE or M2X medium, each containing 1 µg/mL chloramphenicol. These cultures were diluted to an OD_660_ of 0.001 and incubated overnight at 30°C with shaking at 200 rpm. The next morning, these cultures were used to inoculate 500 µL of liquid medium in the wells of a clear 48-well plate. The plate lids were sealed onto the plate body with AeraSeal sealing film (RPI Research Products International) and incubated at 30°C with shaking at 155 RPM. After 1.5 or 2.5 h, OD_660_ and fluorescence (excitation at 497 ± 10 nm; emission at 523 ± 10 nm) were measured on a Tecan Spark 20M plate reader, and fluorescence values were normalized to the corresponding absorbance. Statistical analysis was conducted using GraphPad Prism version 10.1 or R.

### Antibiotic/chemical sensitivity assays

To evaluate the susceptibility of *C. crescentus* strains to cefiderocol, streptonigrin, and CHIR-090, 200 µL of cell suspension adjusted to the same optical density (OD_660_ = 0.3) was spread onto the surface of PYE agar plates. For cefiderocol, an antibiotic gradient strip (Liofilchem MIC test strips) was placed centrally on each plate. For streptonigrin (Sigma-Aldrich) or CHIR-090 (Sigma-Aldrich), a 6-mm filter disc was loaded with 15 µL of 1 mg/mL streptonigrin or 2 mg/mL CHIR-090. The extent of sensitivity was determined by measuring the zone of inhibition.

### Extraction of cell membrane proteins

Strains were grown overnight in 2 mL of PYE at 30°C. The next day, these cultures were used to inoculate into 50 mL of PYE in the evening at an OD_660_ of 0.001. Once cultures reached an OD_660_ 0.4 they were induced with 100 µM dipyridyl and grown for 6 hours. Cultures were pelleted at 7197*g* for 10 minutes and the supernatants were removed. Pellets were then resuspended in 1 mL of ice-cold Tris pH 7.4 with 1 µL of DNAse and 1 µL of RNAse H. The resuspended cultures were lysed through sonication through 3-4 cycles of 20 second intervals with an amplitude of 20% and pulse of 1 second ON and 1 second OFF. Detergent based outer membrane protein extraction was accomplished as described in (67). For centrifugation steps during protein extraction, samples were spun at 20,000*g*.

### SDS-PAGE separation of membrane fractions and protein mass spectrometry

Total protein concentration of 1.5 mg/mL was boiled in SDS sample buffer with 3% β-mercaptoethanol for 5 minutes. The samples were electrophoresed at room temperature in a 7.5% polyacrylamide gel. The gels were stained with Coomassie blue. Gels were destained with acetic acid solution and the bands containing the CciO protein were excised.

Gel bands were digested in-gel according to (88) with modifications. Briefly, gel bands were dehydrated using 100% acetonitrile and incubated with 10mM dithiothreitol in 100mM ammonium bicarbonate, pH∼8, at 56C for 45min, dehydrated again and incubated in the dark with 50mM iodoacetamide in 100mM ammonium bicarbonate for 20min. Gel bands were then washed with ammonium bicarbonate and dehydrated again. Sequencing grade modified trypsin was prepared to 0.005ug/uL in 50mM ammonium bicarbonate and ∼50uL of this was added to each gel band so that the gel was completely submerged. Bands were then incubated at 37C overnight. Peptides were extracted from the gel by water bath sonication in a solution of 60%Acetonitrile (ACN)/1%Trifluoroacetic acid (TFA) and vacuum dried to ∼2uL. Peptides were then re-suspended in 2% ACN/0.1%TFA to 20uL. From this, 5uL were automatically injected by a Thermo (www.thermo.com) EASYnLC 1200 onto a Thermo Acclaim PepMap RSLC 0.1mm x 20mm c18 trapping column and washed for ∼5min with buffer A. Bound peptides were then eluted onto a Thermo Acclaim PepMap RSLC 0.075mm x 500mm C18 analytical column over 35min with a gradient of 8%B to 40%B in 24min, ramping to 90%B at 25min and held at 90%B for the duration of the run (Buffer A = 99.9% Water/0.1% Formic Acid, Buffer B = 80% ACN/0.1% Formic Acid/19.9% Water). Column temperature was maintained at 50C using an integrated column heater (PRSO-V2, www.sonation.com).

Eluted peptides were sprayed into a ThermoFisher Q-Exactive HF-X mass spectrometer (www.thermo.com) using a FlexSpray spray ion source. Survey scans were taken in the Orbi trap (60000 resolution, determined at m/z 200) and the top 15 ions in each survey scan are then subjected to automatic higher energy collision induced dissociation (HCD) with fragment spectra acquired at 15000 resolution. The resulting MS/MS spectra were converted to peak lists using Mascot Distiller, v2.8.1 (www.matrixscience.com) and searched against all *C. crescentus* (*vibrioides*) protein entries available from Uniprot (downloaded from www.uniprot.org on 20201124), appended with common laboratory contaminants (downloaded from www.thegpm.org, cRAP project), using the Mascot searching algorithm, v 2.8.0.1. The Mascot output was then analyzed using Scaffold Q+S, v5.1.2 (www.proteomesoftware.com) to probabilistically validate protein identifications. Assignments validated using the Scaffold 1%FDR confidence filter are considered true.

### Catecholate supplementation of growth media

A mixture of the iron and catecholate solutions was prepared in a 1:10 molar ratio such that the catecholate was 10-fold more abundant than FeCl_3_. This mix was added to M2X medium as the sole iron source to reach a concentration of 1 µM FeCl_3_ and 10 µM catecholate (2,3-dihydroxybenzoic acid, 3,4-dihydroxybenzoic acid, or chlorogenic acid). Once the medium was prepared, it was immediately used for inoculation as described above for a growth curve.

### Cation supplementation in PYE medium containing 300 µM EDTA

To evaluate the effects of cation supplementation on growth in liquid PYE medium containing 300 µM EDTA, the reagents were mixed in the following manner. First, appropriate volumes of EDTA and cation stock solutions were deposited on the opposite sides of a 50-mL tube. Then, liquid PYE medium was added to the tube and shaken. The mixture was dispensed in 2-mL aliquots into glass culture tubes and inoculated as described above for 16-h growth experiments.

### Cultivation of ***C. crescentus*** in lake water

A colony from the WT, Δ*cciO*, or Δ*cciT* strain grown on PYE agar plates was inoculated into liquid PYE medium in triplicate and grown overnight. The next morning, cells were collected by centrifugation and washed three times with 0.2 µM filtered lake water (collected as described above) by serially pelleting and resuspending the cells. After the last wash, the resuspended cells were used to inoculate filtered lake water at a starting OD_660_ of 0.2 and serially diluted 10-fold three times (for a factor of 10^3^). Then 100 µL of the final dilution was used to inoculate 10 mL of desired lake water condition (Huron, Lansing, or EDTA treated). Culture density was assessed then and after 5, 10, 24, and 29 h of growth by determining the number of colony-forming units.

### Measurement of total organic carbon, nitrogen, and sulfate contents of environmental water samples

Lake Lansing and Lake Huron water samples were submitted to EGLE for content analysis of total organic carbon (TOC), nitrogen, sulfate, chloride, conductivity, and alkalinity. TOC content was measured following the persulfate oxidation method 5310C (ASTM Standards). Nitrogen content was measured following the Total Kjeldahl Nitrogen by Semi-Automated Colorimetry approach (EPA method 351.2). Sulfate ion content was measured following the standard test method for sulfate ion in water (ASTM method D516-16). Additional lake analytical methods are provided in Table S5.

### Inductively coupled plasma mass spectrometry (ICP-MS) analysis

All blanks, calibration standards, and ICP-MS samples were prepared using trace-metal-grade nitric acid (70% [w/v], Fisher Chemical, Cat. No. A509P212), ultrapure water (18.2 MΩ·cm at 25°C, Millipore Sigma Milli-Q^®^ IQ7000 Ultrapure Water Purification System), and metal-free polypropylene conical tubes (15 mL and 50 mL, Labcon, Petaluma, CA, USA). Cellular ^31^P (^31^P^16^O), ^56^Fe, ^63^Cu, and ^66^Zn contents of *C. crescentus* WT and mutant strains were measured by ICP-MS. Harvested bacterial cultures were spin-washed with Milli-Q water with or without 10 micromolar EDTA in the metal-free conical tubes. The bacterial cell pellets were dried in a heating block at 80ºC overnight prior to extraction. For liquid samples such as a medium or lake water samples, 100–500 µL of each sample was also dried in the heating block or oven at 80ºC overnight with the conical tube caps loosened. Dried cell pellets and liquid residues were digested in 150 µL of 70% (w/v) nitric acid at 70ºC for 3–4 h or until the pellet was dissolved completely. Following digestion, the final volume of each ICP-MS sample was brought up to 5 mL with Milli-Q water to yield a 3% (w/v) nitric acid matrix. The elemental concentration of all digested samples and calibration standards were determined on a parts per billion (ppb, µg/L) scale using an Agilent 8900 Triple Quadrupole ICP-MS (Agilent, Santa Clara, CA, USA) equipped with an Agilent SPS 4 Autosampler, integrated sample introduction system (ISIS), x-lens, and micromist nebulizer. Daily tuning of the instrument was accomplished using a tuning solution (Agilent, Cat. No. 5185-5959) containing Li, Co, Y, Ce, and Tl. Global tune optimization was based on optimizing intensities for ^7^Li, ^89^Y, and ^205^Tl while minimizing oxides (^140^Ce^16^O/^140^Ce < 1.5%) and doubly charged species (^140^Ce^++^/^140^Ce^+^ < 2%). Following global instrument tuning, gas mode tuning in He KED and O_2_ mode was accomplished using the same tuning solution. In KED mode (using 100% ultra-high-purity [UHP] He, Airgas), intensities for ^59^Co, ^89^Y, and ^205^Tl were maximized while minimizing oxides (^140^Ce^16^O/^140^Ce < 0.5%) and doubly charged species (^140^Ce^++^/^140^Ce^+^ < 1.5%) with short-term relative standard deviations (RSDs) < 3.5%. In O_2_ mode (using 100% UHP O_2_, Airgas) intensities for ^59^Co, ^89^Y, and ^205^Tl were maximized with short-term RSDs < 3.5%. Internal standardization was accomplished inline using the ISIS valve and a 200-ppb internal standard solution prepared in 3% (w/v) trace nitric acid in ultrapure water consisting of Bi, In, ^6^Li, Sc, Tb, and Y (IV-ICPMS-71D, Inorganic Ventures).

The isotopes selected for analysis of media samples were ^31^P (^31^P^16^O), ^56^Fe, ^57^Fe, ^58^Ni, ^59^Co, ^60^Ni, ^63^Cu, ^64^Zn, ^65^Cu, ^66^Zn, ^75^AS, ^78^Se (^78^Se^16^O), ^95^Mo, ^80^Se (^80^Se^16^O), and ^111^Cd with ^6^Li, ^45^Sc, and ^89^Y used for internal standardization. For ICP-MS calibration standard preparation, a 10-ppm stock solution containing As, Ca, Cd, Co, Cr, Cu, Fe, Gd, K, Mg, Mn, Mo, Na, Ni, P, Pt, S, Se, V, and Zn (IV-65024, Inorganic Ventures, Christiansburg, VA, USA) was diluted with 3% (w/v) trace nitric acid in ultrapure water to a final element concentration that covers a range of 0.02–800 ppb. Element concentrations in each sample were derived from a calibration curve generated by Mass Hunter software (Agilent).

## Notes

### Competing Interest Statement

The authors have declared no competing interest.

### Summary of Updates

This version has been revised based on feedback received during peer review.

