## Supplemental Figures for "A co-conserved gene pair supports *Caulobacter* iron homeostasis during chelation stress"

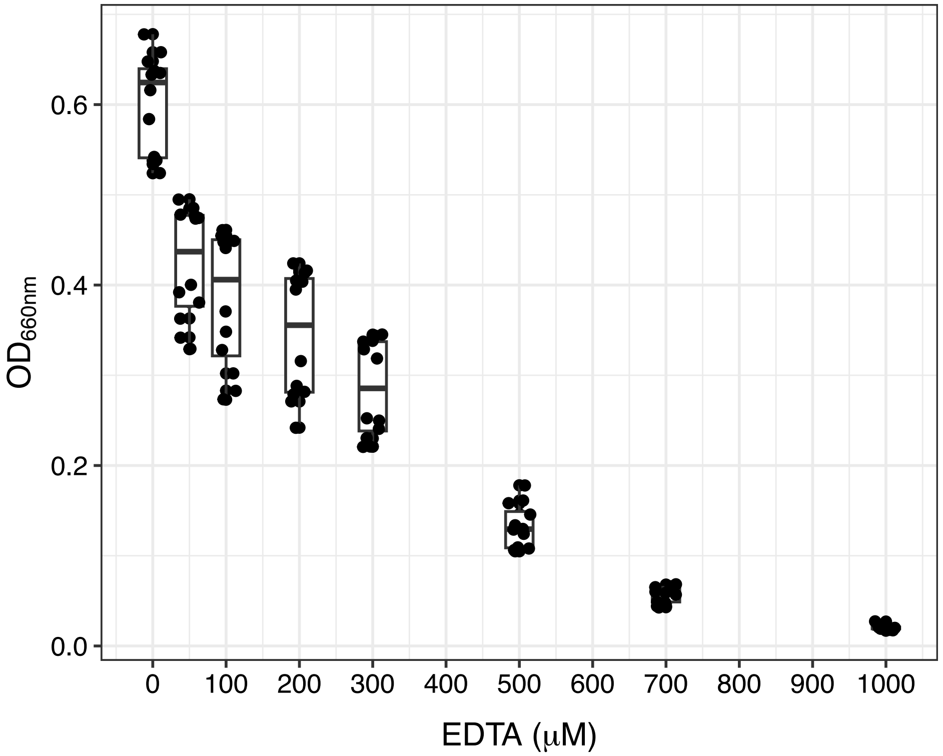


**Figure S1. EDTA attenuates growth of *C. crescentus* in a dose-dependent manner.** Optical density at 660 nm of cultures of *C. crescentus* strain CB15 after 16 h of cultivation in complex liquid medium (PYE) with increasing concentrations of EDTA. All cultures were inoculated at an initial $OD_{660}$ of 0.01. Data represent 12 biological replicates. In the boxplots, the midline represents the median, and the edges of the box represent the 25th and 75th percentiles. Individual data points are shown as black circles.

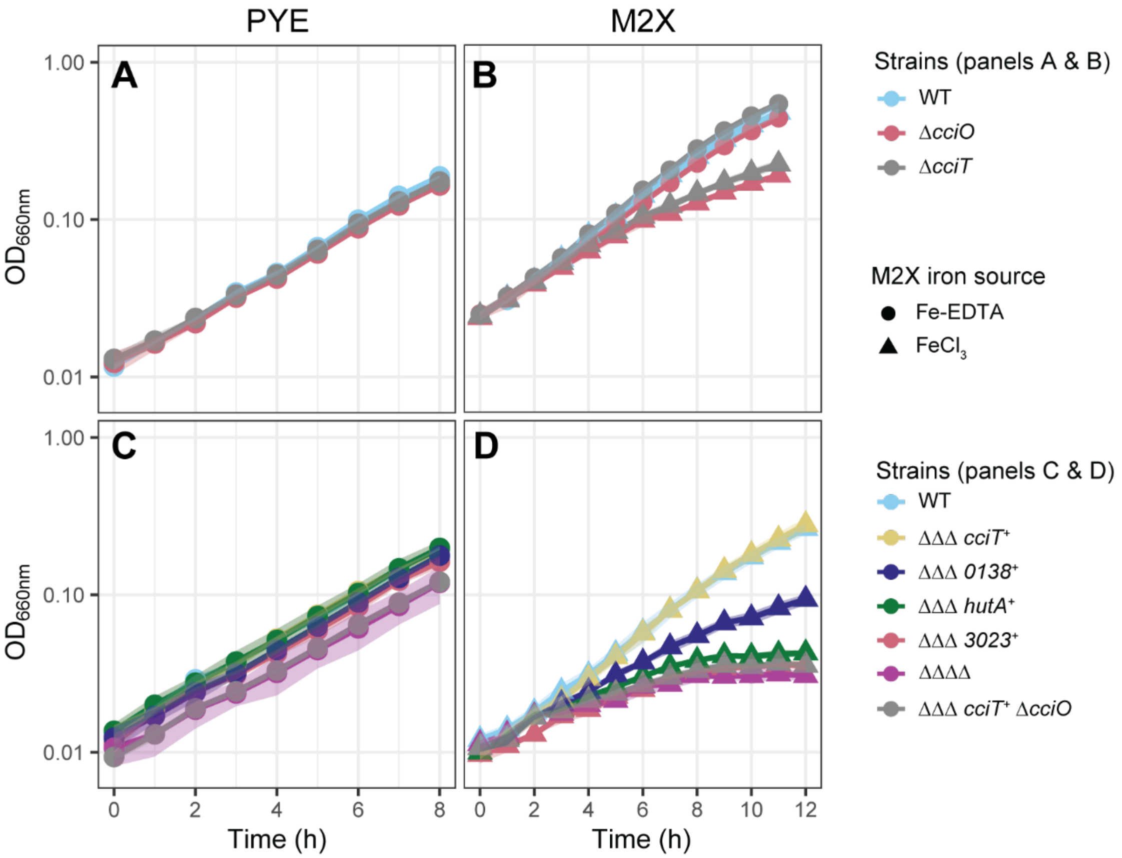


**Figure S2. Growth of *cciT* and *cciO* mutants in complex (PYE) and defined (M2X) media.** (**A&C**) Growth of strains measured by optical density at 660 nm ($OD_{660}$) in complex liquid PYE medium without iron supplementation. (**B&D**) Growth of strains in defined liquid M2X medium where the sole iron source is either 10 μM Fe•EDTA (circles) or 10 μM $FeCl_3$ (triangles). **A&B** show growth of the WT, Δ*cciO*, and Δ*cciT* mutant strains. **C&D** show growth of strains retaining only one of the four Fur-regulated TBDT genes, a strain lacking all four Fur-regulated TBDT genes (ΔΔΔΔ), and a strain in which *cciT* is the only Fur-regulated TBDT gene and *cciO* has been deleted (*cciT*$^+$ ΔΔΔ Δ*cciO*). Values are means ± SD (as shaded areas) from three biological replicates generated on the same day. The SD is too small to be visible in most cases.

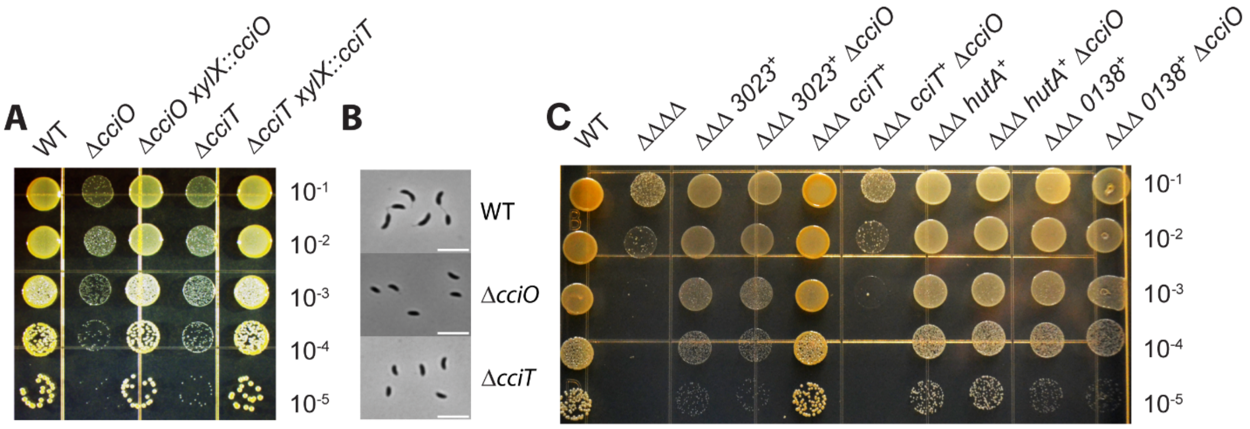


**Figure S3*. cciO* is specifically required for *cciT* to support *C. crescentus* growth on solid medium.** Growth of the same strain set presented in Figure S2 is presented here on solid medium. **(A)** Serial dilutions (10-fold) of WT, Δ*cciO* and Δ*cciT* mutants*,* and genetically complemented strains expressing *cciO* or *cciT* from the *xylX* chromosomal locus. **(B)** Representative micrographs of cells used to determine the cell area distributions of WT, Δ*cciO* and Δ*cciT* in Figure 3. White scale bar in the lower right corner of each micrograph represents 5 µm. **(C)** Serial dilutions of WT and mutant strains retaining only one of the four Fur-regulated TBDT genes (*cciT*, CCNA_00138, CCNA_03023, and *hutA*) in a background encoding or lacking *cciO* (Δ*cciO*). Strains were spotted onto solid complex medium (PYE). Dilution factor is indicated on the right.

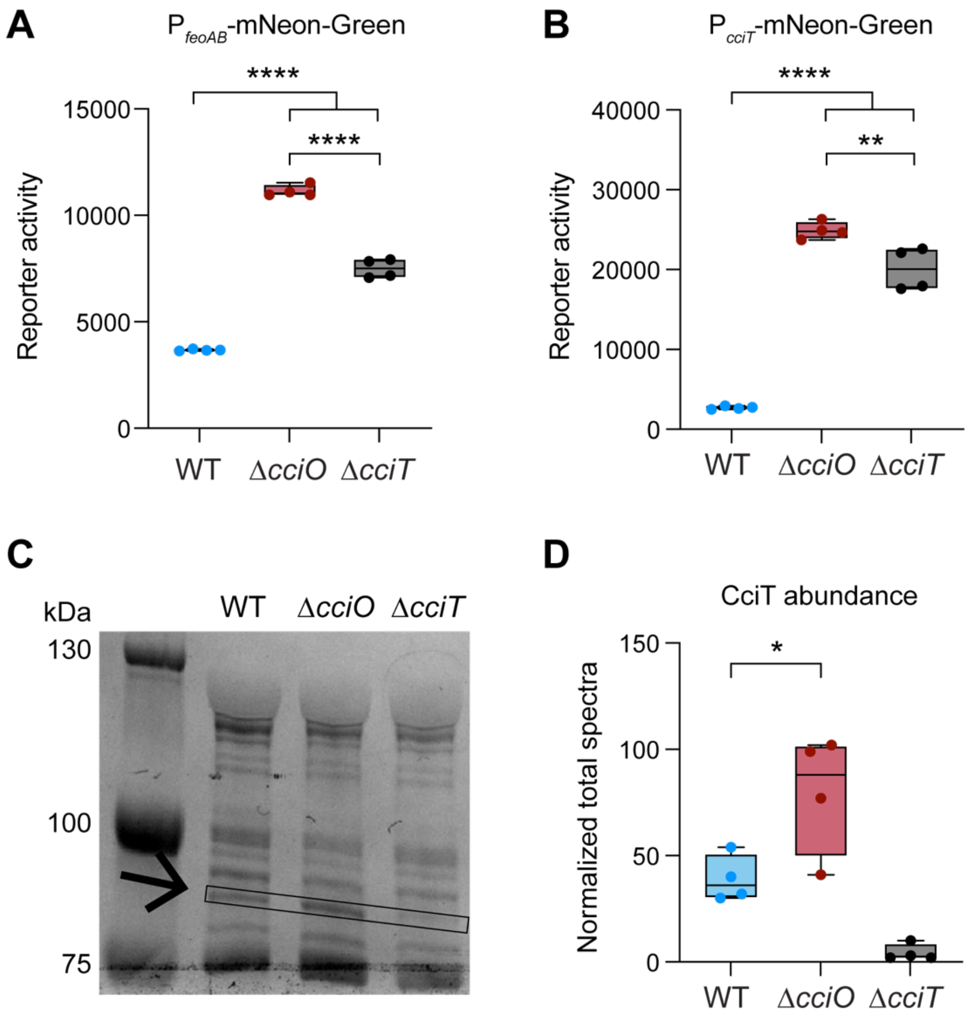


**Figure S4. Transcription reporter and protein mass spectrometry validation of differences in Fur-dependent gene expression in the *cciT* and *cciO* deletion strains.** Fluorescence signal from promoter:mNeonGreen transcriptional reporter cassettes with **(A)** the Fur-regulated *feoAB* promoter ($P_{feoAB}$) or **(B)** the Fur-regulated *cciT* promoter ($P_{cciT}$) in WT, Δ*cciT*, and Δ*cciO* strains grown in the same conditions as in the RNA-seq experiment (M2X defined medium with $FeCl_3$; see Figure 4). **(C)** SDS-PAGE gel of outer membrane protein preparations from WT, Δ*cciT*, and Δ*cciO* strains cultivated in liquid PYE medium containing 100 μM of the intracellular iron chelator dipyridyl. Membrane proteins were extracted as described (67). The band corresponding to the size of CciT is marked with a box and arrow on the Coomassie brilliant blue-stained gel. **(D)** Mass spectrometry quantification of CciT tryptic peptides isolated from gel slices cut from the marked box in panel C. Peptides were quantified from four independent outer-membrane protein preparations. The significance of differences in transcription or tryptic peptide levels was determined by one-way ANOVA followed by Tukey's post-hoc test for each pairwise comparison except in D, where we only compared CciT levels in WT and Δ*cciO* because the transport protein is not produced in Δ*cciT* (* $p < 0.05$; ** $p < 0.01$; **** $p < 0.00001$).

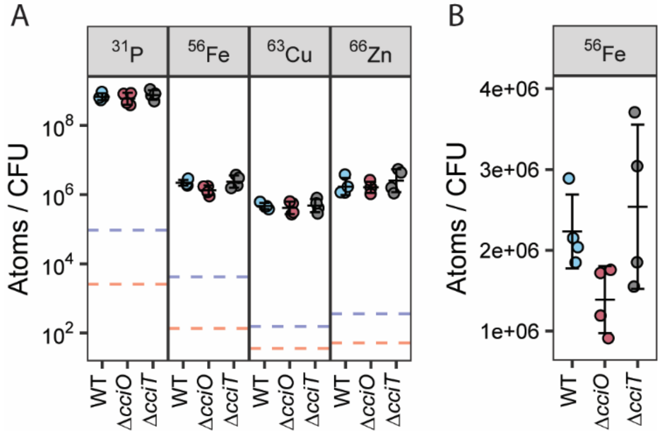


**Figure S5. (A)** ICP-QQQ-MS bio-elemental analysis of 4 biological replicates of WT, Δ*cciO*, and Δ*cciT* strains of *C. crescentus* cultivated in peptone-yeast extract (PYE) complex medium. Phosphorus (P), iron (Fe), copper (Cu), and zinc (Zn) concentrations for each replicate normalized to the number of colony-forming units. The dashed blue line represents the background equivalent concentration (BEC) of the element in the digestion blanks. The dashed orange line represents the detection limit (DL) of the element. The data points shown have been corrected for BEC during the calibration process. The BEC and DL lines are provided to demonstrate the instrument's detection capability and the background matrix concentration. **(B)** Zoomed view of Fe data (atoms/CFU) for WT, Δ*cciO*, and Δ*cciT* strains.

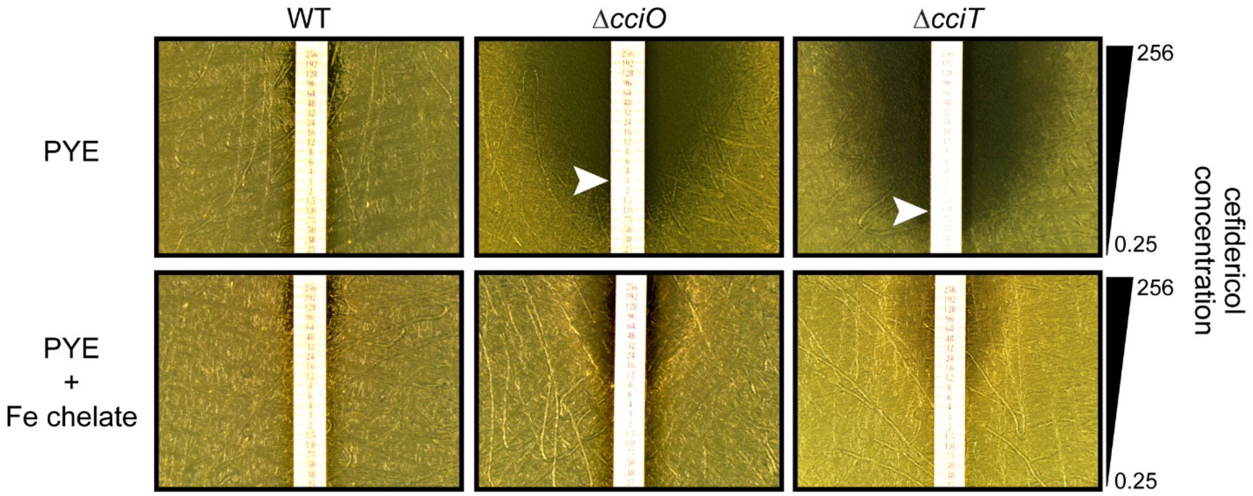


**Figure S6.** Sensitivity of *C. crescentus* wild-type (WT) and strains lacking *cciO* or *cciT* to the catechol-cephalosporin “Trojan horse” antibiotic, cefiderocol. Tests were conducted on complex (PYE) medium without (top) and with (bottom) supplementation of the growth medium with 10 μM Fe•EDTA. Arrowheads indicate approximate minimal inhibitory concentration in each condition. Lack of arrow indicates resistance to highest concentration of cefiderocol on the strip. The range of cefiderocol concentration in the strip is indicated on the right.

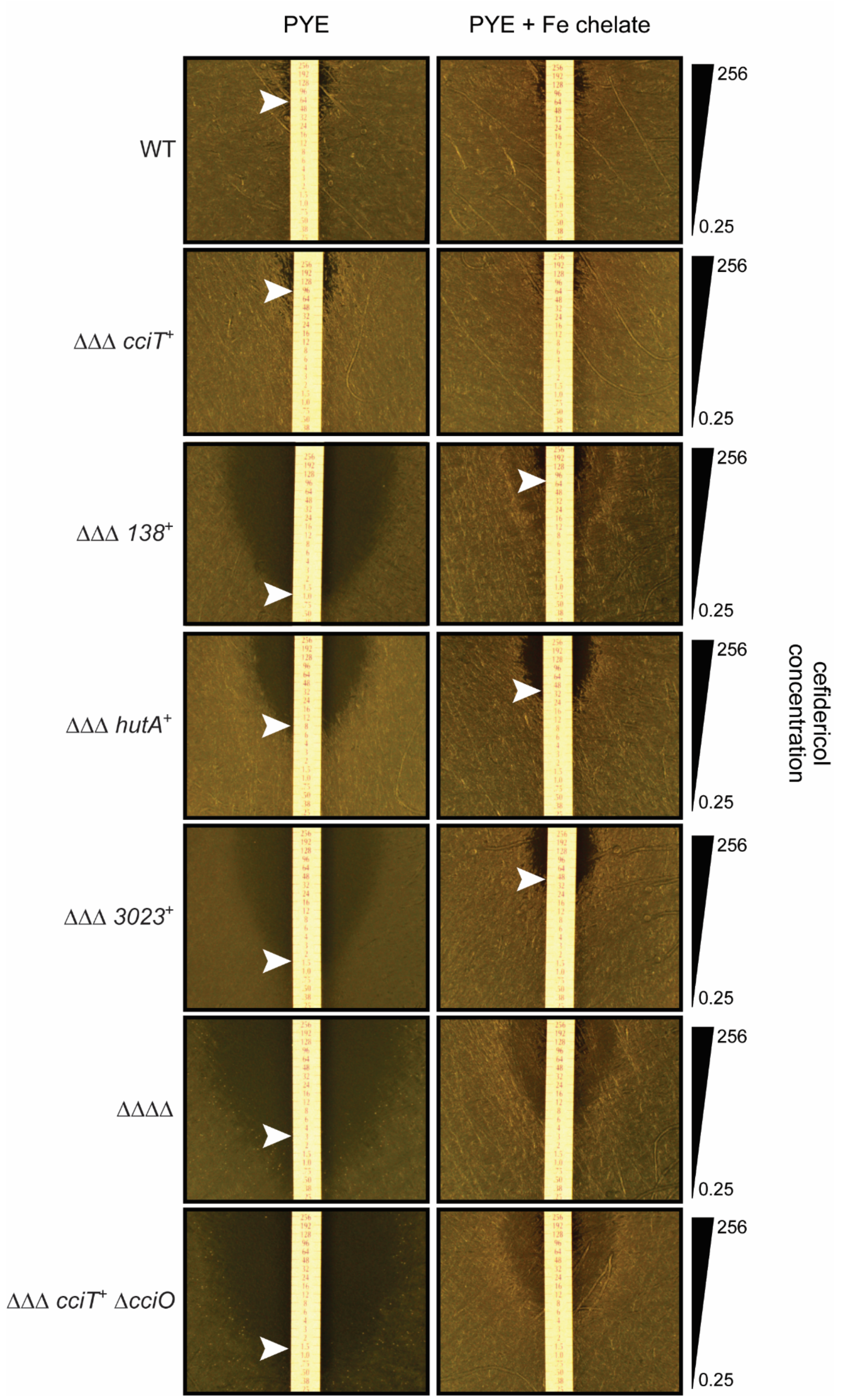


**Figure S7.** Wild-type (WT) *C. crescentus* and a strain expressing *cciT* but lacking the other three Fur-regulated TBDT genes (*cciT*$^{+}$ ΔΔΔ) are resistant to cefiderocol. Any strain lacking *cciO or cciT* is sensitized to cefiderocol. Addition of 10 μM Fe•EDTA to the medium both improved growth and conferred some cefiderocol resistance compared to that in PYE medium alone. Minimal inhibitory concentration is marked with a white arrow. Lack of arrow indicates resistance to highest concentration of cefiderocol on the strip. The range of cefiderocol concentration in the strip is indicated on the right

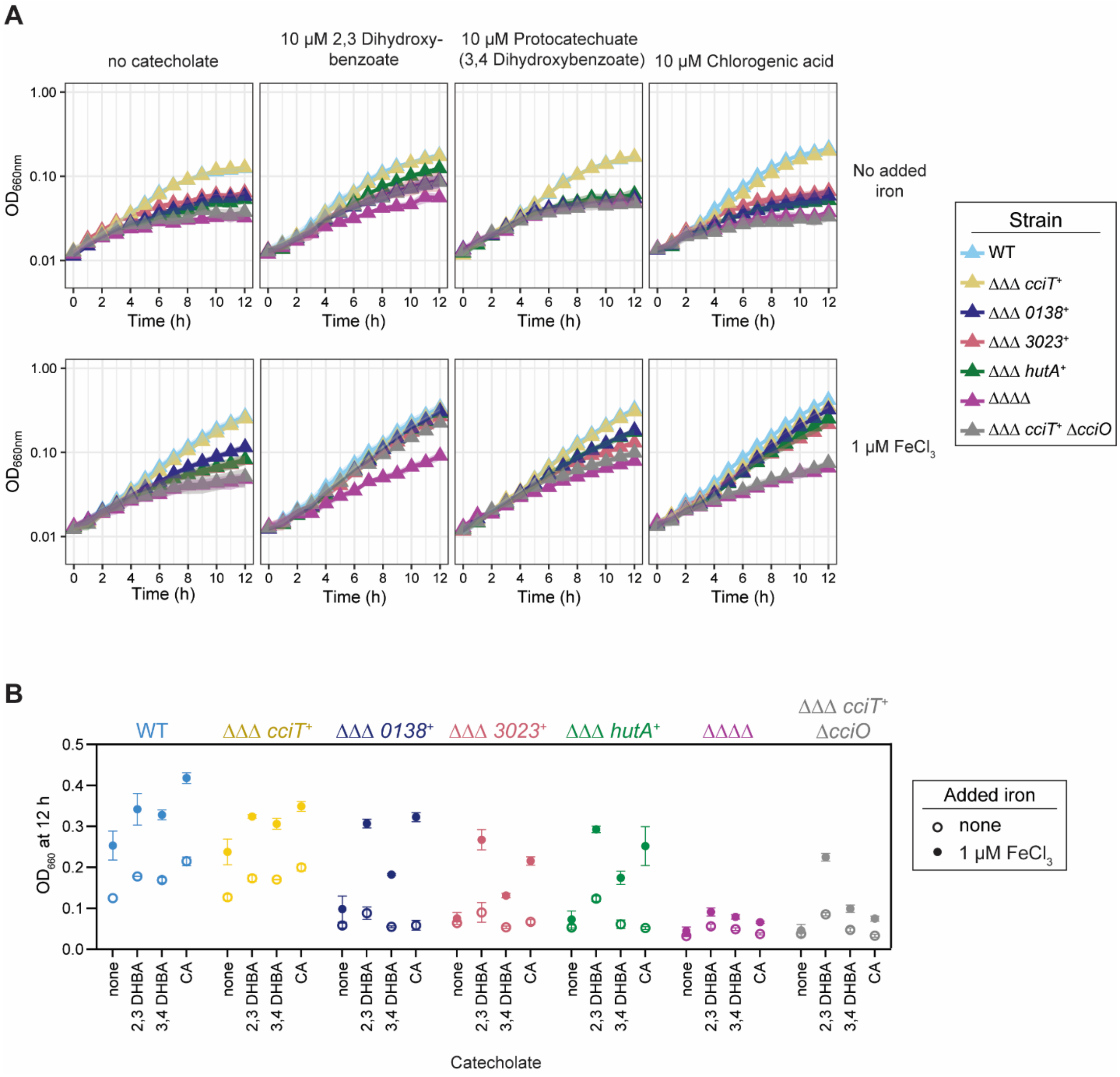


**Figure S8. Effects of catecholates and iron on growth in defined medium. (A)** Growth curves of strains grown in defined liquid M2X-based medium with no added iron (top) or with the addition of 1 μM $FeCl_3$ (bottom), in the presence of 10 μM of three different catecholates (labeled above). **(B)** Optical densities at 660 nm of cultures after 12 h of growth, extracted from the curves in A.

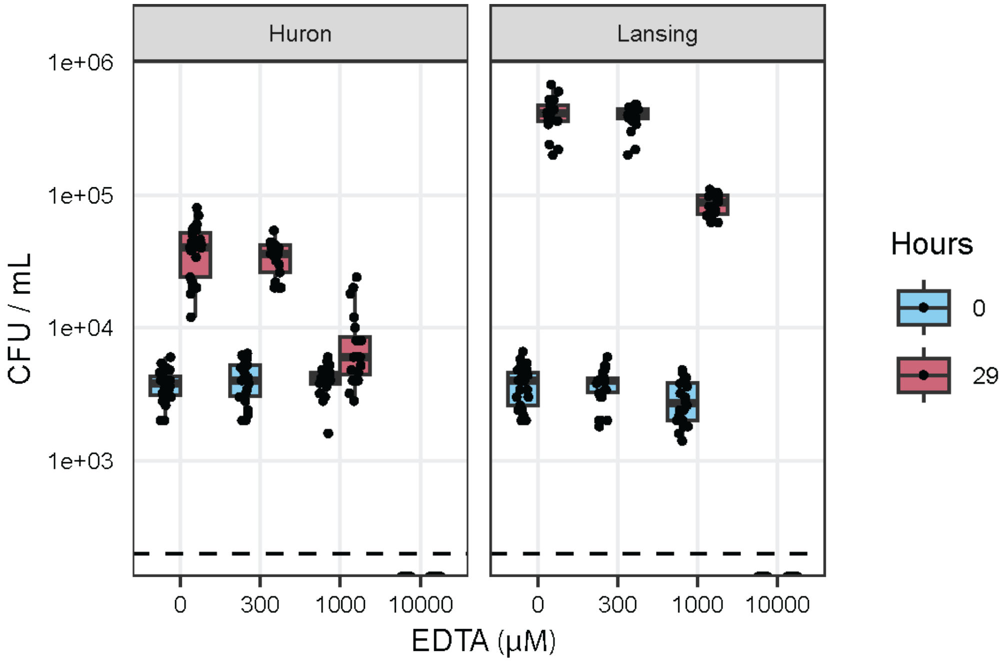


**Figure S9. Determination of EDTA toxicity to wild-type *C. crescentus* in Lake Huron and Lake Lansing water.** Wild-type (WT) *C. crecsentus* was cultivated from a starting titer of ~3900 CFU/ml in water from Lakes Lansing and Huron supplemented with increasing concentrations of EDTA. Cell densities were measured by enumeration of colony forming units (CFU) immediately after EDTA treatment (0 hours; blue) and after 29 hours (red) of growth. Summary box plots represent the median, 25$^{th}$ and 75$^{th}$ percentiles. Dashed line indicates limit of detection (200 CFU/ml).

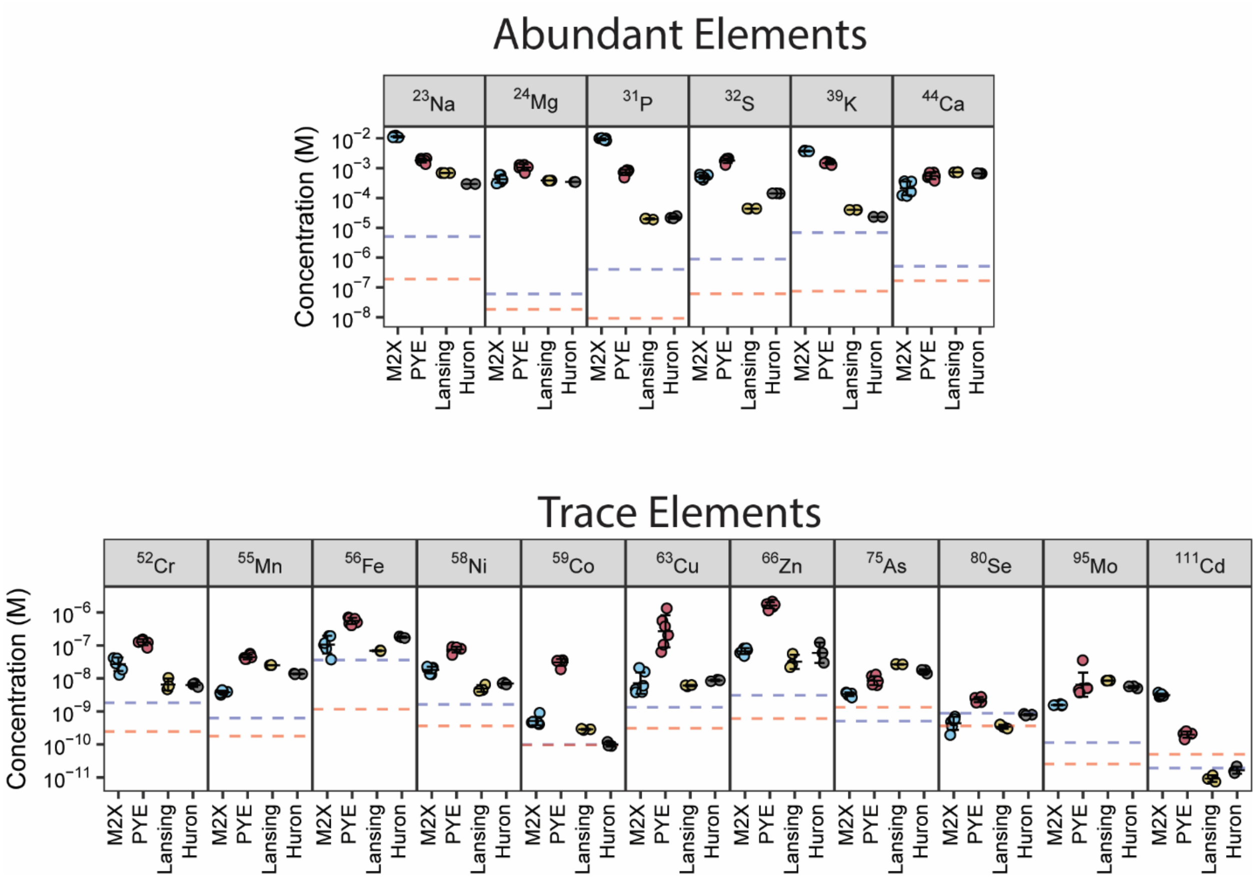


**Figure S10. ICP-QQQ-MS elemental analysis of growth media.** Quantitative elemental profiles of liquid M2X medium (without added iron supplement), liquid PYE medium, and water collected from Hammond Bay in Lake Huron (collected November 2023) and Lake Lansing public dock at South Lake Lansing Park (collected February 2024), measured by ICP-MS-QQQ. Data represent nine independent measurements for each condition (3 independent samples measured 3 times). Mean ± SD are overlayed with the individual data points. The laboratory media were prepared three times and measured in technical triplicates. The lake sources were each measured on one sampling day where three samples were prepared and measured in technical triplicate. The dashed blue line represents the background equivalent concentration (BEC) of the element in the digestion blanks. The dashed orange line represents the detection limit (DL) of the element. The data points shown have been corrected for BEC during the calibration process. The BEC and DL lines are provided to demonstrate the instrument's detection capability and the background matrix concentration.

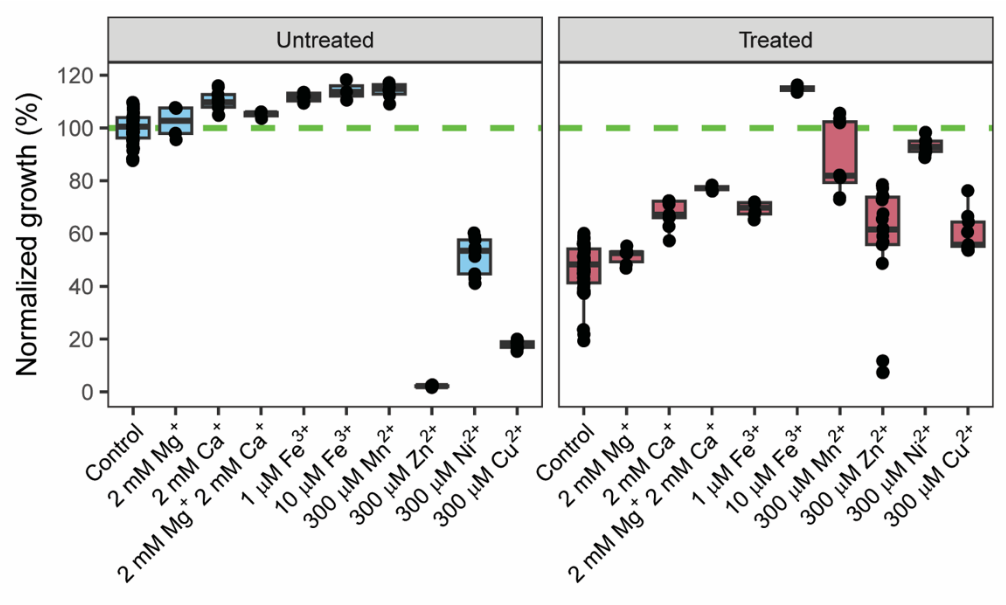


**Figure S11. Rescue of EDTA-dependent growth inhibition by select cations.** Density of *C. crescentus* cultures after 16 h of growth of was measured spectrophotometrically ($OD_{660}$) in the presence of indicated cations, without (Untreated) or with (Treated) 300 µM EDTA. Values were normalized to WT growth in untreated (Control) PYE medium. For each condition, at least three biological replicates were measured. Summary box plots representing the median, 25$^{th}$ and 75$^{th}$ percentiles are overlayed with the individual data points.
